# Gut microbe-derived lactic acid optimizes host energy metabolism during starvation

**DOI:** 10.1101/2025.05.27.656452

**Authors:** Jason William Millington, Jamie Alcira Lopez, Amin M. Sajjadian, Robert J. Scheffler, Brian C. DeFelice, William Basil Ludington, Benjamin H. Good, Lucy Erin O’Brien, Kerwyn Casey Huang

## Abstract

Gut microbes convert dietary compounds into an array of metabolites that can directly provide energy to their host and indirectly impact host metabolism as systemic endocrine signals. Here, we show that gut microbe-derived metabolites can extend *Drosophila melanogaster* survival during starvation, despite minimal alteration of dietary energy intake. Combining survival assays with mathematical modeling and untargeted metabolomics, we identify a single, dominant mediator of starvation resilience: lactic acid produced by the commensal bacterium *Lactiplantibacillus plantarum*. We discover that the basis of starvation resilience is not catabolism of lactic acid using lactate dehydrogenase, but rather increased dietary energy yield through lactic acid-driven promotion of oxidative phosphorylation. Our findings emphasize the role of the microbiome as a source of endocrine cues coordinating host metabolism and underscore the potential of microbiome-derived metabolites as therapeutic molecules for manipulating metabolic health and preventing disease.

## Introduction

Gut microbes have emerged as a key regulator of host energy metabolism^1–4^. In response to host dietary intake, they utilize ingested nutrients for growth, increasing microbial biomass and producing an array of diverse metabolites as byproducts. These processes result in many nutrient-driven host-microbe interactions, including competition for nutrient resources, cross-feeding of microbe-derived metabolites that are taken up by the host to generate energy, and production of signaling molecules that coordinate changes to host metabolism in a manner evocative of endocrine regulation through hormones^5–8^. Each aspect of these interactions has important consequences for regulating host energy metabolism, and intestinal microbiome dysbiosis has been associated with metabolic disease^9^.

An extreme example of adaptive regulation of energy metabolism is starvation resilience^10^. Metabolic flexibility to endure unpredictable cycles of feast-famine is a desirable trait for many animal species in the wild, and persistent global prevalence of human malnourishment highlights its relevance to public health. Intriguingly, gut microbes have been successfully deployed as probiotics to treat malnourishment in animal models^11–17^. However, the underlying mechanisms through which microbes promote starvation resilience remain largely unknown.

The fruit fly, *Drosophila melanogaster*, is a well-established model for the study of energy metabolism regulation during starvation^18,19^, particularly with regards to host-microbe interactions. In *Drosophila*, *Lactiplantibacillus* and *Acetobacter* species dominate the microbiome^20–24^ and have been implicated in regulating host energy metabolism and lifespan^13–15,25–31^. Microbiome composition correlates with starvation resilience in *Drosophila*^32–34^, potentially occurring via the landscape of microbe-derived metabolites that may act directly as nutrients or indirectly as endocrine signals. However, although a few candidate microbe-derived metabolites have been proposed to impact host energy metabolism^35–37^, we lack systematic approaches to identify microbe-derived regulators of energy metabolism. Thus, the immense, yet poorly characterized, chemical diversity produced by the microbiome provides a rich ground for discovering regulation mechanisms.

Here, we develop an experimental platform and mathematical modeling framework to systematically interrogate the impact of molecules on *Drosophila* energy metabolism during starvation. We apply this method to the metabolites produced by *Drosophila* gut microbiome commensals and find that these metabolites can substantially extend lifespan during starvation despite minimal alteration to dietary caloric content. We identify that *Lactiplantibacillus plantarum*-derived lactic acid is necessary and sufficient for starvation survival extension, and show that lactic acid extends survival via modulation of dietary energy yield in a manner largely independent of the metabolic conversion of lactic acid to pyruvate. We hypothesize that this mechanism acts instead via lactic acid-mediated regulation of oxidative phosphorylation. Consistent with our hypothesis, we show that pharmacological inhibition of oxidative phosphorylation abolishes the lactic acid-mediated starvation resilience. By demonstrating that a single microbial metabolite globally remodels host energy metabolism for starvation resilience, our results highlight potential roles of the microbiome as a source of endocrine signals coordinating metabolic regulation.

## Results

### Microbe-derived metabolites prolong Drosophila starvation survival

To investigate how nutrient-based host-microbe interactions contribute to host energy metabolism in *Drosophila*, we developed an assay to screen for altered host starvation survival based on feeding flies microbial spent media (Fig. 1A). In flies near starvation, we found that gut microbiome load and composition were significantly altered relative to fed conditions (Fig. S1A-C). To amplify the host impact of microbe-derived metabolites, we utilized a chemically defined medium (CDM) designed for growth of fly gut microbes^38^. CDM captures two key aspects necessary for our screening approach: (1) CDM permits the growth of all fly gut commensal bacteria and results in high levels of microbial metabolite production, and (2) CDM is a near-starvation baseline condition for fly survival. By itself, 1X CDM supports a marginal fly lifespan of ∼1 week (Fig. 1C), versus 2-3 days in complete starvation (STV) conditions and ∼2-3 months under *ad libitum* conditions^39^. To control for potential media batch effects, all assays were performed with the relevant controls including STV and/or CDM conditions.

**Figure 1:**
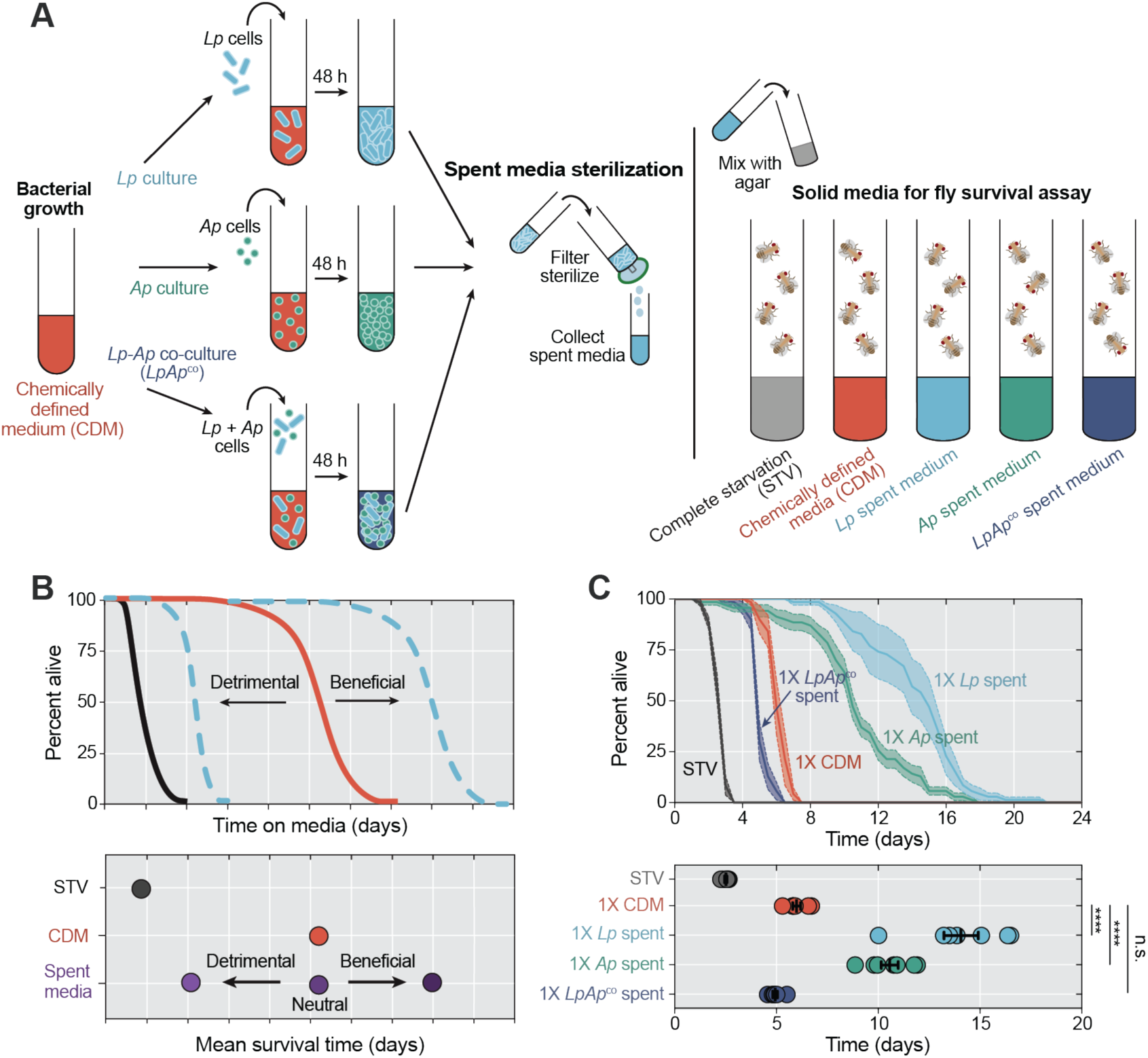
Microbe-derived metabolites impact *Drosophila* starvation survival. A) Schematic of *Drosophila* survival assay in flies fed chemically defined media (CDM) supernatant versus bacterial spent media and expected outcomes and interpretations. Spent media from stationary phase (48 h) cultures were obtained from fly gut commensals (*Lp* and *Ap*) grown in monoculture and co-culture by centrifugation and filtering before mixing with agar to make a solid substrate for fly survival quantification. B) Survival is determined by nutrient-driven interactions impacting energy yield, with the potential for both increased survival from beneficial metabolites and shortened survival from e.g., nutrient competition. C) Mean survival was significantly higher in *Ore-R* mated females fed *Lp-* and *Ap*-spent media compared to CDM (*p* < 0.0001 for both comparisons; one-way ANOVA followed by Dunnett’s multiple comparison test). *N* = 7 biological replicates, *n* = 70 flies per condition. Mean survival was not significantly different between *Ore-R* mated females fed *LpAp*^co^*-*spent medium compared to flies fed CDM supernatant (*p* = 0.3; one-way ANOVA followed by Dunnett’s multiple comparison test). *N* = 7 biological replicates, *n* = 70 flies per condition. ****: *p* < 0.0001; ns: not significant; error bars indicate 1 SEM.

We used this assay to characterize the contribution to host starvation survival of two abundant *Drosophila* gut commensals isolated from wild flies: *Lactiplantibacillus plantarum* (hereafter *Lp*) and *Acetobacter pasteurianus* (*Ap*)^31,40^. By filtering stationary phase bacteria cultures, we generated *Lp*- and *Ap*-derived cell-free spent media, from which we produced solid substrates that we fed to adult wild-type *Oregon-R* mated female flies and quantified survival (Fig. 1B,C). Our null expectation was that host-microbe resource competition would result in reduced fly survival on spent media compared to CDM alone due to bacterial consumption of resources that would otherwise have been available to the fly. Indeed, flies fed the spent medium from an *LpAp* coculture (*LpAp*^co^) had a mean survival time of 5 days, slightly shorter than on fresh CDM (Fig. 1C). However, we found that flies fed 1X spent medium from either *Lp* or *Ap* lived significantly longer than those fed fresh CDM (mean survival of 14 days and 10.5 days, respectively, versus 6 days on CDM; Fig. 1C), indicating that microbe-derived metabolites can positively impact starvation resilience. *Lp-* and *Ap-*spent media prolonged survival without altering fly food intake or taste preference (Fig. S2A-D), active energy expenditure (Fig. S2E-G), or egg production (Fig. S2H). Altogether, these results argue against changes in food intake or active energy expenditure mediating prolonged survival and instead suggest that other aspects of energy metabolism are impacted by microbe-derived metabolites.

### Energy balance models suggest that microbe-derived metabolites prolong survival beyond increasing dietary energy content

A multitude of factors could be responsible for the observed lifespan extension in spent media, including changes in nutritional content, fly behavior, and/or fly metabolism. To disentangle these potential mechanisms mediating differential survival, we utilized mathematical models to interpret survival data through a framework based on changes in energy balance. This framework considers organisms from a thermodynamic perspective and has been widely applied to starvation and body mass dynamics across vertebrate and invertebrate species^41–46^. In these models, survival during starvation is determined by the initial amount of stored energy and the rates of energy intake and expenditure. Once energy stores are consumed, organismal death follows (Fig. 2A). Since the observed distributions of survival times were unimodal, we focused on the mean and variance of fly survival times across feeding conditions.

**Figure 2:**
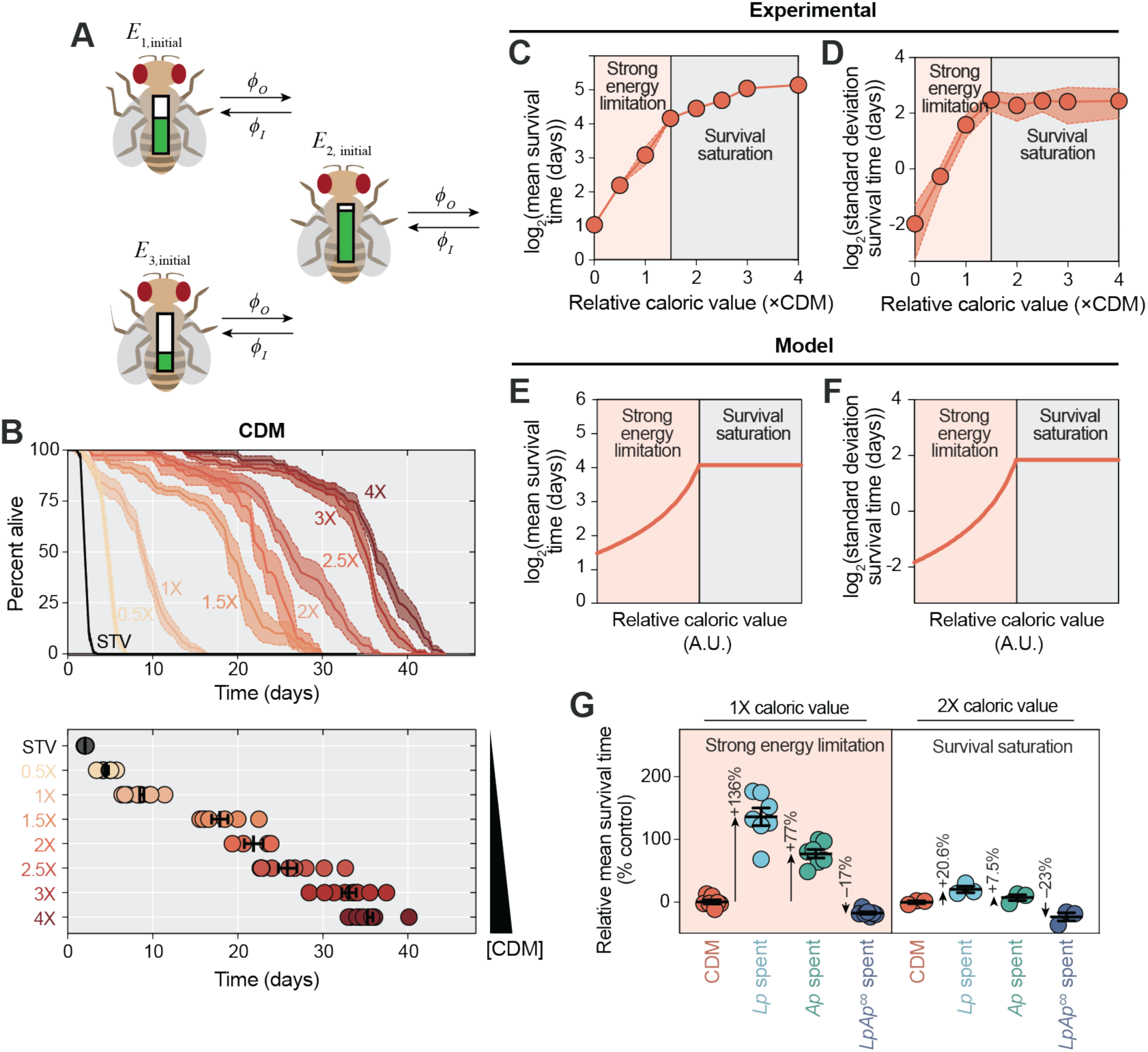
Microbe-derived metabolites cause survival to deviate from simple energy balance models in starvation regimes with substantial energy limitation. A) Schematic of energy balance model used to interpret *Drosophila* survival data as described by Eq. 1. This model considers a population of flies with mean initial energy storage *μ*_*E*_ and each fly expends energy at flux *Ø*_O_ and intakes energy at flux *Ø*_*I*_. B) Mean survival increased with increasing CDM concentration. C,D) Mean survival (C) and variance (D) increased superlinearly between 0-1.5X CDM, and saturate between 2-4X CDM. E,F) Energy balance model predictions for mean survival (E) and variance (F) as a function of increasing relative caloric value. Survival is predicted to be superlinear in the energy limitation regime and to saturate when energy demands for normal lifespan are met. G) The percent difference in mean survival relative to the control was higher at lower caloric values than at high caloric values for *Lp*- and *Ap*-spent media. Error bars indicate 1 SEM.

Our model considers a population of flies, each with a time-dependent total energy store *E*_*i*_(*t*) such that death occurs deterministically when *E*_*i*_ = 0. For simplicity, we assume that variability in fly survival emerges from random variation across the population in the initial value of *E*_*i*_at the time starvation commences (*t*=0), which we assume has mean *μ*_*E*_ and variance 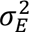. Each fly expends energy at rate *Ø*_O_ and intakes energy at a rate *Ø*_*I*_, which is dependent on the energy extracted from the fly diet. Starvation commences when intake drops below expenditure (*Ø*_*I*_ < *Ø*_O_). For simplicity, we assume here that these rates are constant across the population and over time for a given diet (see Methods for an analysis of variable rates in energy input/output across the population).

The mean and standard deviation of survival times is dictated by the mean and standard deviation in initial values of *E*_*i*_, scaled by 1/(*Ø*_O_ − *Ø*_*I*_) (Methods):

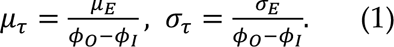

When the fly is near starvation (*Ø*_*I*_∼*Ø*_O_), small changes in dietary energy extraction or energy expenditure can lead to superlinear sensitivity of survival times to energy metabolism changes (Methods). The similar scaling of *μ*_*τ*_ and *σ*_*τ*_ with respect to (*Ø*_*I*_ − *Ø*_O_) provides two complementary tests for the model. Note that Eq. 1 predicts a non-physical infinite lifespan when *Ø*_O_ = *Ø*_*I*_; in practice, lifespan will saturate in nutrient-replete conditions, eventually leading to a reduction in sensitivity to energy metabolism changes (Methods).

We first sought to test whether it could successfully predict the quantitative dependence of fly survival dynamics on caloric intake on fresh CDM. We varied caloric intake (*Ø*_*I*_) by feeding flies CDM at concentrations ranging from 0.5X-4X, which resulted in two regimes of behavior for the mean and standard deviation of survival time (*μ*_*τ*_ and *σ*_*τ*_, respectively): between 0.5X-1.5X CDM (low *Ø*_*I*_, the energy limitation regime), *μ*_*τ*_ and *σ*_*τ*_ increased superlinearly with increasing caloric value (Fig. 2B-D). For 2X-4X CDM (high *Ø*_*I*_), *μ*_*τ*_ and *σ*_*τ*_ saturated (Fig. 2C,D), presumably because energy demands are largely rescued by nutrients present in the CDM (Fig. 2B). These data indicate that the transition to the saturated regime occurs above 1X CDM concentration, hence 1X CDM is a useful baseline condition due to the high lifespan sensitivity around this level of energy intake. Moreover, the agreement between our data and the predictions of our energy balance model across varying caloric intake for low *Ø*_*I*_ (Fig. 2E,F) suggest that fly survival on low levels of CDM is driven by simple energy balance, establishing a baseline from which to interrogate the mechanisms underlying extended survival in spent-media conditions.

We next asked whether the observed effects of spent media (Fig. 1C) are consistent with straightforward changes in caloric value. As microbes grow in CDM, nutrients are consumed and converted to biomass and microbe-derived metabolites. While we do not know the identity of all microbe-derived metabolites, we can infer the approximate change in the energy available to the fly. Most of the energy content of CDM (as estimated by the Gibbs free energy of combustion) comes from glucose and amino acids, both of which are accessible to the fly (Supplementary Text). Hence, it is unlikely that microbial metabolism increases survival by increasing energy content; in fact, as microbes grow, they utilize some of the energy present in the media (Supplementary Text), lowering the total available energy in the media relative to fresh CDM. Thus, the observation that spent media from *Lp* and *Ap* monocultures extend fly survival rather than shorten it (Fig. 1C) suggests that microbe-derived metabolites alter the way *Drosophila* extracts, expends, or stores energy. Based on the expectations of our energy balance model, we concluded that the small observed differences in activity, food intake volume, and egg production are quantitatively insufficient to explain the lifespan extension in flies fed *Lp*-spent media (Fig. S2I,J).

Next, we sought to determine whether our energy balance model captures the survival dynamics of flies fed spent media from *Lp*, *Ap*, or *LpAp*^co^ at different initial caloric values. Our analysis of CDM alone suggests that if survival is indeed controlled by energy balance, *Lp-* and *Ap-*spent media should have minimal effects relative to CDM alone at higher concentrations in the saturated regime. To test this prediction, we measured fly survival at 2X caloric density for flies fed CDM, or *Lp-*, *Ap-*, or *LpAp*^co^-spent media. Flies fed 2X *Lp-* and *Ap-*spent media had a mean survival time of ∼26 days and 24 days, respectively. These survival dynamics are only marginally longer than the ∼23 day mean survival of flies fed 2X CDM (Fig. 2G, S3), suggesting that the effect of microbe-derived metabolites on survival is strongest in the starvation regime. However, flies fed 2X *LpAp*^co^ spent medium had a significantly shorter mean survival time of ∼17 days compared with CDM-fed flies (Fig. 2G, S3), equivalent to between 1X and 1.5X CDM (Fig. 2B). This observation suggests that the co-culture depletes the media of fly-accessible nutrients; the larger effect relative to 1X *LpAp*^co^ spent medium (Fig. 1C) highlights the superlinearity of survival in the starvation regime below 1.5X relative caloric value (Fig. 2C). Thus, by applying energy balance models, we find that changes in energy metabolism (and not changes in fly behaviors such as food intake) likely underly the starvation resilience of flies fed *Lp*-spent medium.

### Prolonged starvation survival is mediated by Lp-derived lactic acid

We next sought to identify the microbe-derived metabolite(s) that promote fly starvation survival. To this end, we analyzed the chemical content of CDM and of *Lp*-, *Ap*-, and *LpAp*^co^-spent media via liquid chromatography with tandem mass spectrometry (LC-MS, Fig. 3A). We identified ∼1500 metabolomic features post-filtering (mostly unannotated), with distinct metabolomic profiles across media (Fig. 3B, Methods). One enriched metabolite in *Lp*-spent medium was lactic acid, as expected given the metabolic capabilities of *Lp*, which was undetectable in CDM and *Ap*-spent medium (Fig. 3C). Lactic acid was still present in *LpAp*^co^-spent medium, but at significantly lower abundance (Fig. 3C). While other metabolites were also enriched in spent media, we initially focused on lactic acid as it is the primary metabolic product of *Lp* fermentation and *Lp*-spent medium provided the largest lifespan extension (Fig. 1C).

**Figure 3:**
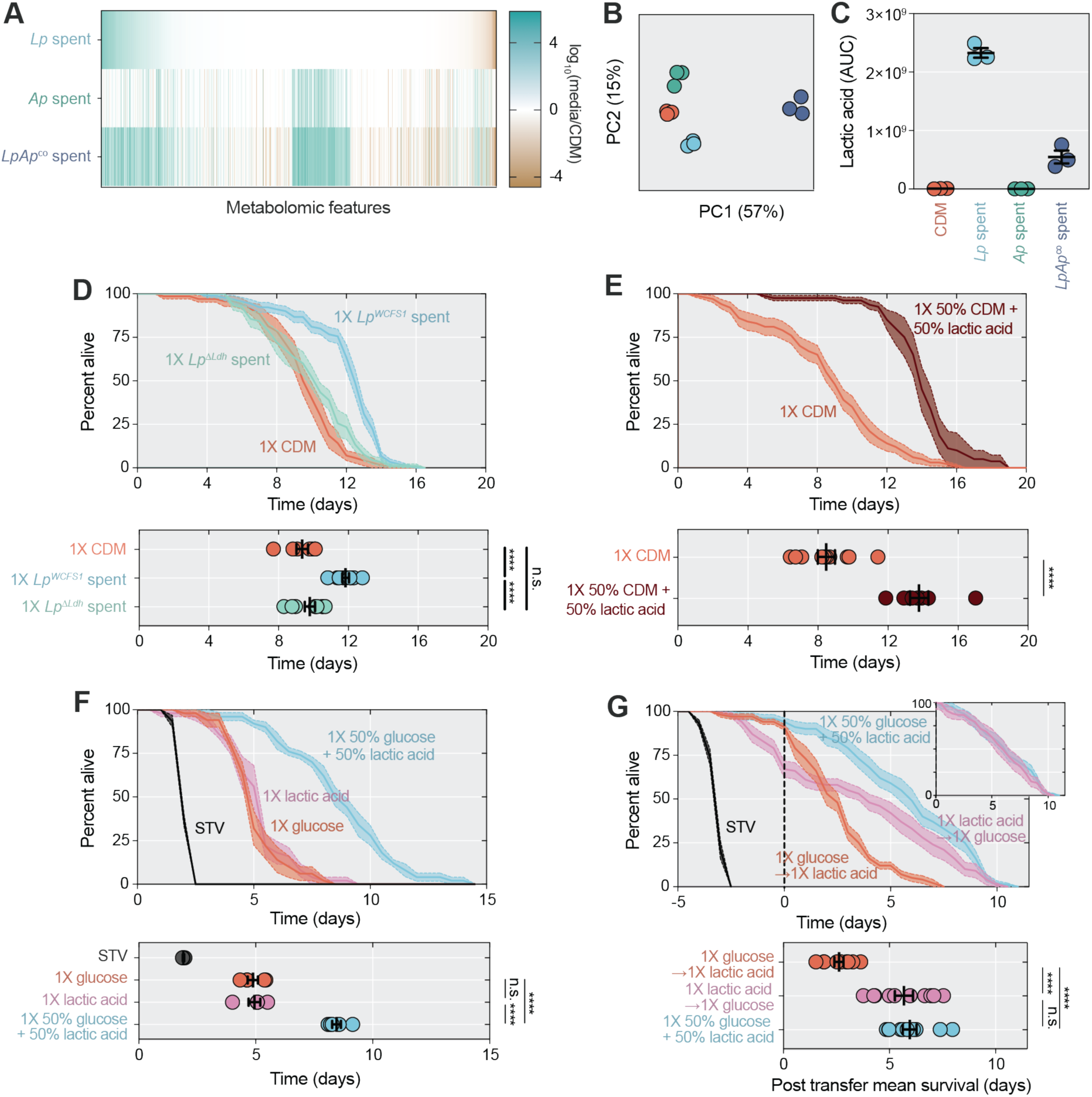
*Lp*-derived lactic acid is necessary and sufficient for starvation survival and acts via an interaction with glucose metabolism. A) Mean fold change of LC-MS signal intensity from spent media relative to fresh CDM supernatant for each species across all metabolomic features, annotated or otherwise. *N*=3 biological replicates. B) Principal component analysis of fresh CDM supernatant and spent media from monocultures and co-culture. C) Lactic acid levels in fresh CDM supernatant and spent media from monocultures and co-culture. Lactic acid was significantly more concentrated in *Lp*-compared with *LpAp*^co^-spent medium (*p* = 0.0002; one-way ANOVA followed by Tukey HSD test). *N* = 3 biological replicates. D) Mean survival was significantly lower in *Ore-R* mated females fed spent medium from the *Lp*^Δ*Ldh*^ mutant or CDM supernatant compared with flies fed spent medium from the parent strain *Lp^WCFS1^* (*p* < 0.0001 for both comparisons; one-way ANOVA followed by Tukey HSD test). Mean survival was not significantly different between *Ore-R* mated females fed *Lp*^Δ*Ldh*^-spent medium compared to flies fed CDM supernatant (*p* = 0.51; one-way ANOVA followed by Tukey HSD test). *N* = 7-9 biological replicates, *n* = 70-90 flies per condition. E) Mean survival was significantly higher in *Ore-R* mated females fed 1X 50% CDM + 50% lactic acid compared to flies fed 1X CDM (*p* < 0.0001; Student’s *t* test). *N* = 8-10 biological replicates, *n* = 80-100 flies per condition. F) Mean survival was significantly higher in *Ore-R* mated females fed 1X 50% glucose + 50% lactic acid compared to flies fed 1X glucose or 1X lactic acid (*p* < 0.0001 for both comparisons; one-way ANOVA followed by Tukey HSD test). Mean survival was not significantly different between *Ore-R* mated females fed 1X glucose compared to flies fed 1X lactic acid (*p* = 0.98; one-way ANOVA followed by Tukey HSD test). *N* = 5 biological replicates, *n* = 50 flies per condition. G) Mean survival was significantly higher in *Ore-R* mated females fed 1X lactic acid for 5 days and then switched to 1X glucose after 5 days of lactic acid feeding compared to flies with the opposite feeding regimen (1X glucose followed by 1X lactic acid) or flies maintained on 1X 50% glucose + 50% lactic acid mixture (*p* < 0.0001 for both comparisons; one-way ANOVA followed by Tukey HSD test). Mean survival was not significantly different between *Ore-R* mated females first fed 1X lactic acid followed by 1X glucose compared to flies maintained on the 1X 50% glucose + 50% lactic acid mixture (*p* = 0.83; one-way ANOVA followed by Tukey HSD test). *N* = 10 biological replicates, *n* = 100 flies per condition. ****: *p* < 0.0001; ns: not significant; error bars indicate 1 SEM.

Given the pronounced enrichment of lactic acid in *Lp*-spent medium, we sought to determine the importance of *Lp*-derived lactic acid on fly starvation survival. We utilized *Lp*^Δ^*^Ldh^*, a strain incapable of lactic acid production due to deletion of lactate dehydrogenase (*Ldh*), which catalyzes the terminal fermentation of pyruvate to produce lactic acid. Flies fed *Lp*^Δ^*^Ldh^*-spent medium had significantly shorter survival times than spent medium from the parent *Lp^WCFS1^* strain (Fig. 3D) and similar to that of CDM-fed controls. This finding establishes *Lp*-derived lactic acid as necessary for increased starvation survival, and that *Lp*^Δ^*^Ldh^* does not substantially deplete CDM of nutrients that benefit fly survival.

To examine whether supplementation of fresh CDM with lactic acid alone could extend starvation survival, we diluted CDM two-fold and added DL-lactic acid (hereafter, “lactic acid”) to create near-isocaloric, lactic acid-supplemented CDM (0.5X CDM + 62.5 mM lactic acid). We found that lactic acid supplementation was sufficient to significantly extend starvation survival compared to CDM-fed flies (Fig. 3E). As with *Lp*-spent media, the effects of lactic acid were dependent on caloric value: at twice the calories (1X CDM + 125 mM lactic acid), survival was unaffected compared to 2X CDM (Fig. S4A,B). Thus, lactic acid is sufficient to extend survival in the starvation regime but has no effect at higher caloric values as lifespan saturates. We further investigated whether the deprotonated form of lactate (conjugate base) was sufficient for this effect.

We found directly feeding lactate provided no nutritional value, consistent with our prediction that lactic acid dominates in spent media (Fig. S5A-D). However, as we do not know which form predominates intracellularly and thus do not rule out a role for intracellular lactate mediating this effect.

### Lactic acid prolongs starvation survival through an interaction with glucose metabolism

Given that *Lp*-spent medium extends starvation survival in a manner dependent on lactic acid but independent of alterations to food intake and active energy expenditure (Fig. S2D,E), we next considered alternate mechanisms of lactic acid action. One possibility is a lactic acid-dependent increase in energy extraction from dietary or stored nutrients (effectively increasing *Ø*_*I*_ or *μ*_*E*_). To determine the relative bioavailability of lactic acid for energy production, we fed flies isocaloric solutions of either lactic acid or glucose (the main carbon source in CDM). If lactic acid is more bioavailable than glucose, then the lactic acid diet should promote longer starvation survival. However, such an effect was not observed experimentally: isocaloric diets of glucose (1X D-glucose, hereafter glucose) or lactic acid (1X lactic acid) led to similar survival lifespans (Fig. S4C,D). Furthermore, doubling the concentration of lactic acid (2X lactic acid) had no effect on starvation survival compared to 1X lactic acid (Fig. S4C,D). On the other hand, higher concentrations of glucose increased starvation survival (Fig. S4C,D), consistent with an energy balance mechanism. These findings exclude increased bioavailability of lactic acid as the mechanism through which it prolongs starvation survival.

An alternative hypothesis is that lactic acid prolongs starvation survival by interacting with metabolism of other nutrients such as glucose. Supporting this possibility, we found that flies fed a 1:1 mixture of glucose and lactic acid at 1X caloric value survived significantly longer than flies fed either 1X glucose or 1X lactic acid alone (Fig. 3F; S4C,D). Since all three diets were isocaloric, the survival extension produced by the combined mixture suggests that lactic acid in the presence of glucose enables increased dietary energy extraction.

Given the lack of support for a direct energy balance mechanism, we next considered a signaling role for lactic acid that impacts energy metabolism. This hypothesis is consistent with known links between lactate and signaling through G-protein coupled receptor-81 (GPR81) that impacts lipolysis and energy metabolism in mammals^47,48^. In contrast to nutrients, which are utilized rapidly to impact energy metabolism, signaling molecules can mediate long-term metabolic effects. We hypothesized that if lactic acid has a signaling role, then priming via an initial interval of lactic acid feeding could potentiate increased survival during subsequent feeding with glucose. To test this hypothesis, we fed flies lactic acid for 5 days before transferring them to an isocaloric glucose diet (without lactic acid) on which flies were maintained until death. We also performed the reciprocal treatment of 5 days of glucose feeding followed by transfer to lactic acid, as well as a control condition of flies maintained on a mixture of glucose and lactic acid. We found that flies fed lactic acid prior to glucose feeding survived significantly longer than flies transferred from glucose to lactic acid (Fig. 3G). Moreover, the survival dynamics post-transfer from lactic acid to glucose were quantitatively similar to those for the glucose and lactic acid mixture (Fig. 3G). These findings suggest that lactic acid has a priming effect on glucose metabolism that extends starvation survival even after lactic acid is removed, consistent with a signaling role for lactic acid.

### Prolonged starvation survival does not require lactic acid breakdown

If lactic acid impacts starvation survival via signaling, we expected that this effect would be independent of lactic acid metabolism. Bacterial Ldh (Fig. 3D), which is functionally and structurally divergent from metazoan Ldh, nearly exclusively produces lactate from pyruvate. In contrast, in metazoans, lactate is metabolized to pyruvate in a reversible reaction catalyzed by Ldh, regenerating NAD^+^ in the process^49^. Pyruvate is then shuttled to the mitochondria to generate ATP and fuel further glycolysis^50^. To determine the importance of host lactic acid metabolism, we examined the effect of lactic acid on starvation survival in flies whose *Ldh* activity was reduced through ubiquitous, RNAi-mediated knockdown of *Ldh* in adulthood (*tubulin-GAL4, tubulin-GAL80^ts^*). Flies were fed isocaloric media containing either glucose, lactic acid, or a mixture of glucose and lactic acid, and their survival was compared to control flies. If *Ldh* activity is necessary for the mixture of glucose and lactic acid to prolong survival, then we expected *Ldh-RNAi* flies to survive for a shorter interval than control flies.

However, we found that in both the knockdown and control genotypes, the mixture of glucose and lactic acid prolonged survival compared to glucose only (Fig. S6A-C). The lactic acid-only medium also prolonged survival of *Ldh-RNAi* flies but not control flies, suggesting RNAi-specific effects that may be related to a decrease in lactic acid-induced stress. Together, these data suggest that Ldh is not limiting for lactic acid to promote starvation survival in the presence of glucose, consistent with a ignaling role for lactic acid.

### Lactic acid delays stored energy utilization

How can a signaling mechanism facilitate increased survival without higher food intake or reduced energy expenditure? Given our priming data, we hypothesized that lactic acid mediates an increase in food energy yield; specifically, that ingestion of glucose and lactic acid together allows the fly to extract more energy than ingestion of only lactic acid or only glucose alone. In this case, feeding with glucose and lactic acid together should allow flies to mobilize their stores of body fat more slowly. To test this prediction, we investigated the rate of stored body fat breakdown by measuring levels of whole-body triglycerides, the main energy storage molecule in *Drosophila* and other higher organisms^51,52^. During complete nutrient starvation, flies exhibited rapid triglyceride mobilization (Fig. 4A), as expected. However, flies fed a mixture of glucose and lactic acid delayed their fat mobilization relative to flies fed only glucose or lactic acid (Fig. 4A,B). These data are consistent with a mechanism in which lactic acid mediates an increase in dietary energy yield.

**Figure 4:**
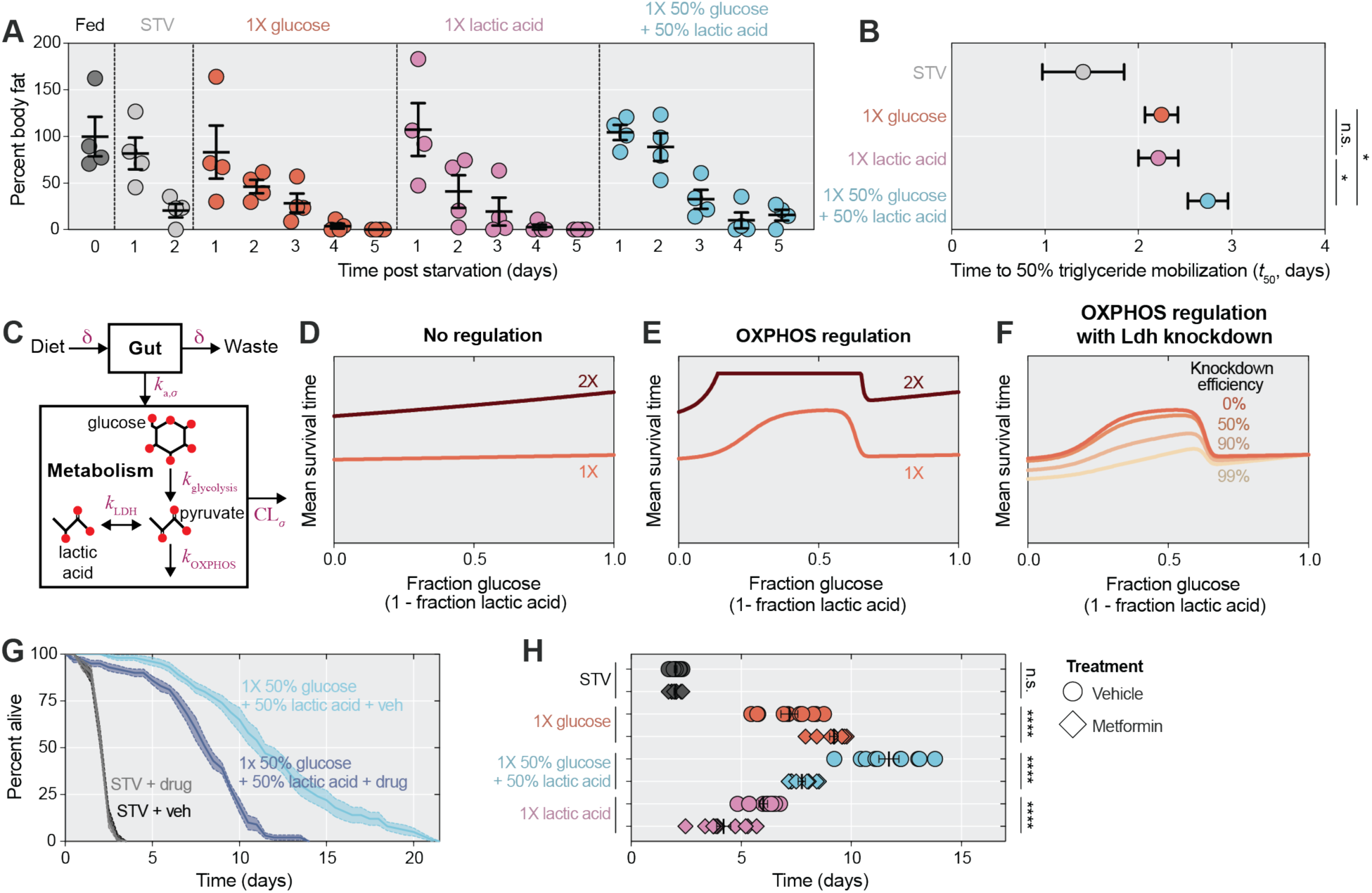
Lactic acid impacts starvation survival via regulation of oxidative phosphorylation (OXPHOS). A) Percentage body fat in the form of triglycerides over 5 days of starvation in *Ore-R* mated females fed 1X glucose, 1X lactic acid, or 1X 50% glucose + 50% lactic acid. *N* = 4 biological replicates, *n* = 20 flies per condition. B) The rate of triglyceride mobilization as determined by a linear fit was significantly slower in flies fed 1X 50% glucose + 50% lactic acid compared to flies fed 1X glucose or 1X lactic acid (*p* = 0.017 and 0.012, respectively; one-way ANOVA followed by Tukey HSD test). The rate of triglyceride mobilization was not significantly different between *Ore-R* mated females fed 1X glucose compared to flies fed 1X lactic acid (*p* = 0.97; one-way ANOVA followed by Tukey HSD test). C) Schematic of pharmacokinetic-pharmacodynamic (PKPD) model used to interpret survival data. This model considers the fate of glucose and lactic acid in *Drosophila* using a simplified representation of central carbon metabolism pathways. D) In the absence of regulation, PKPD model predicts equivalent mean survival between flies fed any ratio of glucose and lactic acid at either low or high caloric values. E) PKPD model with regulation of OXPHOS dependent on the ratio of lactic acid to glucose levels predicts increased mean survival at intermediate ratios. The increase in survival is highest at low caloric values. F) PKPD model with OXPHOS regulation predicts increased mean survival across a range of Ldh knockdown efficiencies. The magnitude of increased survival due to lactic acid addition to a glucose diet is highest with full Ldh expression. G,H) Mean survival was significantly lower in *Ore-R* mated females treated with 25 mM metformin and fed 1X 50% glucose + 50% lactic acid compared to vehicle-treated flies with the same diet (*p* < 0.0001; two-way ANOVA followed by Sidak’s multiple comparison test). Metformin treatment did not affect mean survival for complete starvation in *Ore-R* mated females (*p* > 0.99; two-way ANOVA followed by Sidak’s multiple comparison test). Mean survival was significantly higher in *Ore-R* mated females treated with 25 mM metformin and fed 1X glucose compared to vehicle-treated flies fed 1X glucose (*p* < 0.0001; two-way ANOVA followed by Sidak’s multiple comparison test). Mean survival was significantly lower in *Ore-R* mated females treated with 25 mM metformin and fed 1X lactic acid compared to vehicle-treated flies fed 1X lactic acid (*p* < 0.0001; two-way ANOVA followed by Sidak’s multiple comparison test). *N* = 10 biological replicates, *n* = 100 flies per condition. *: *p* < 0.05, ****: *p* < 0.0001; ns: not significant; error bars indicate 1 SEM.

### Lactic acid prolongs starvation survival by potentiating OXPHOS

In the context of our energy balance model, increased energy yield corresponds to an effective increase in *Ø*_*I*_. To generate hypotheses regarding the mechanisms responsible for such an increase, we sought to generate the simplest model of *Drosophila* energy metabolism consistent with three observations: (1) a mixture of lactic acid and glucose promotes longer survival than glucose or lactic acid alone, (2) the survival benefit of the mixture saturates with increasing caloric content, and (3) the survival benefit of the mixture is largely host Ldh independent. Additionally, the model should exhibit regimes in which glucose and lactic acid alone provide similar survival benefits (Fig. 3F). We focused on pharmacokinetic-pharmacodynamic (PKPD) models, a framework with established efficacy for predicting metabolic outputs and the impact of effector molecules in complex organisms^53^.

Our model (Methods) considers two compartments: the gut and the rest of the body cavity (referred to as the central compartment) (Fig. 4C). The diet contains a mixture of glucose and lactic acid, which flows through the gut and is absorbed into the central compartment. Once absorbed, components undergo a highly simplified set of energy metabolism reactions. Glucose undergoes glycolysis, producing 2 pyruvate and 2 ATP. Lactic acid is reversibly converted to pyruvate, and each pyruvate can be processed via oxidative phosphorylation (hereafter OXPHOS), which produces 17 ATP. Each component is cleared from the central compartment, by excretion from the body or other component-specific metabolic reactions. To connect this PKPD model to the energy balance model, we assume that energy extraction *Ø*_*I*_ from the diet is proportional to ATP production.

We found that in the absence of active regulation of glycolysis and OXPHOS, the dietary energy flux *Ø*_*I*_ is predicted by the PK model to be a linear function of dietary lactic acid and glucose concentrations. Thus, this form of the model does not capture the nonlinear behavior we observed (Fig. 4D). We arrived at similar conclusions using a genome-scale model of *Drosophila* metabolism that considers ∼8,000 reactions and factors such as redox balance (Fig S9A; Methods). These findings are consistent with our empirical results implying a signaling mechanism whereby lactic acid modulates energy metabolism.

To evaluate whether signaling mechanisms could account for our key experimental results, we focused on regulation of OXPHOS, which is where the most energy can be harvested and impacts both lactic acid and glucose energy yield. OXPHOS regulation is also in line with a previous *in vitro* study showing that lactic acid can promote oxidative phosphorylation in HepG2 cells and allow a higher yield of energy from glucose^54^.

Indeed, we found that a regulatory scheme in which the ratio of gut lactic acid to glucose regulates OXPHOS can recapitulate all key behaviors if (1) there is a basal level of OXPHOS activity in lactic acid or glucose diets, and (2) there is an optimal ratio of lactic acid to glucose (Fig. 4E). Furthermore, simulations across increasing levels of Ldh knockdown efficiency produced similar results (Fig. 4F), indicating that an OXPHOS regulatory scheme can be largely Ldh independent. Our proposed regulatory scheme is consistent with several possible mechanisms, such as gene expression altering OXPHOS enzyme levels or post-translational modification of OXPHOS enzymes. Nonmonotonic changes in the per-pyruvate ATP yield of OXPHOS would produce similar effects. Other nonlinear OXPHOS regulatory schemes, such as an optimal concentration of lactic acid rather than an optimal ratio of lactic acid to glucose concentrations, are also potentially consistent with our results.

This model predicts that survival of flies fed a mixture of lactic acid and glucose would be adversely impacted by inhibition of OXPHOS. To test this prediction, we inhibited OXPHOS pharmacologically using metformin, which inhibits OXPHOS by blocking complex I of the electron transport chain^55^, and assessed the impact on starvation survival. In vehicle-treated control flies, we observed that flies fed a mixture of glucose and lactic acid survived significantly longer than glucose or lactic acid fed flies alone (Fig. 4G,H; S7). In contrast, flies fed the mixture in conjunction with metformin treatment did not survive longer than glucose-fed flies alone and their survival was significantly shorter than vehicle-treated controls (Fig. 4G,H; S7). Our observation that metformin abolishes extended survival in flies fed the mixture of glucose and lactic acid supports the model that lactic acid impacts starvation survival via OXPHOS regulation.

## Discussion

Here, we showed that under starvation conditions, lactic acid generated by the gut commensal *Lp* extends host survival by enhancing dietary energy yield via OXPHOS. By developing and applying a method for screening and interrogating the impact of candidate effectors on energy metabolism in *Drosophila*, we uncovered a class of host-microbiome interactions in which a microbial metabolite alters host energy metabolism by modulating the energy yield from nutrient sources. We propose that the gut microbiome, a rich source of metabolic effectors, effectively serves as a cross-kingdom endocrine regulator of host energy metabolism.

Our experiments involved diets pre-conditioned by microbes, a context in which the microbes initially have exclusive access to dietary components. While this pre-conditioning mimics microbially fermented foods, it is unclear to what extent gut microbes chemically modify dietary components *in situ*. As a case study, we examined the lactic acid flux produced by *Lp* in non-starving *Drosophila*. By combining data on fly gut bacterial population sizes, gut transit times, and food intake measurements, we calculated that the flux of lactic acid produced by the fly gut microbiome is similar in magnitude to the flux of carbon compounds from food intake (Supplementary Text).

This suggests that *in situ* bacterial metabolism can indeed generate physiologically relevant fluxes of effector metabolites. Note that owing to the smaller bacterial populations in starved flies (Fig. S1), these fluxes may be less relevant in severe starvation conditions. Our calculation is also consistent with results from other multicellular organisms. In humans, it is well-established that microbial metabolism of dietary compounds can produce physiologically relevant fluxes of fermentation products, supplying up to 10% of incoming energy^35^. Indeed, work on microbiome drug metabolism has shown that microbial modification of even non-food ingested compounds can substantially impact host physiology in mice^56,57^ and humans^58^. Thus, *in situ* chemical modification of ingested compounds may influence organismal physiology across the tree of life.

Our results also reveal a counter-intuitive relationship between dietary energy yield and organismal fitness. The role of diet in human energy balance can be largely captured by assuming that the energy yield is near the thermodynamic limit (i.e., the free energy of combustion)^59,60^ and humans rarely excrete non-CO_2_ carbonaceous waste^50^. However, we found that even under starvation, flies did not always fully utilize the energy contained in their diet, with flies fed 1X CDM leaving at least ∼45% of the dietary energy unutilized (calculated from the increase in energy intake required for *Lp-* mediated survival benefit; Fig. 1C). This finding contrasts with naïve expectations for behavior during starvation, when an organism might be expected to be especially energetically efficient to prolong survival. Why then does *Drosophila* underutilize dietary energy during starvation? One possibility is a trade-off between energy yield and other fitness-relevant phenotypes. For example, in single-celled organisms such as bacteria, a trade-off between energy yield and overall energy influx rate has been well characterized^61^, whereby it can be optimal to only partially oxidize substrates to increase total substrate flux and overall energy flux. Similar trade-offs have also been observed in cancer^62^ and yeast cells^63^. However, it is unlikely that this trade-off between substrate flux and yield explains the behavior of *Drosophila* since food intake volume (Fig. S2D), and thus overall substrate flux, does not change between diets. Other forms of trade-offs, such as with fertility, could underlie this behavior. For example, prolonged periods of starvation can lead to infertility in *Drosophila*^64,65^, decoupling lifespan extension during starvation from reproductive fitness. In addition to nutrient yield trade-offs, an alternative explanation is that the unmodified CDM diet is far from typical *Drosophila* diets and that this under-yielding does not often occur in nature. The diet of wild *Drosophila* consists largely of rotten fruit rich in *Lactiplantibacilli*, *Acetobacter*, and yeasts amongst a host of other microbial species producing microbial metabolites, including lactic acid. Thus, a fly in the wild may typically encounter mixtures of glucose and lactic acid in its food source, rather than the pure glucose found in CDM. Further characterization of phenotypes in relation to energy yield across hosts could shed light on these fundamental evolutionary questions.

Finally, while our study focuses on one interaction between lactic acid and glucose metabolism, the experimental and mathematical tools we developed represent a broad platform for detecting and dissecting the role of energy metabolism effectors more generally. The metagenome of intestinal microbial communities encodes a vast potential for microbe-derived metabolite production^66^, many of which can be detected in host circulation^67,68^ and have potential (largely uncharacterized) roles as signaling molecules, akin to the complex hormonal milieu produced by different cell types in an endocrine organ. Our findings highlight the scope for discovery of novel metabolic effectors among microbe-derived metabolites, potentially paralleling the discovery of hormone-based treatments for metabolic disease from mechanistic studies of circulating regulators of energy metabolism such as GLP-1 and analogs such as semaglutide^69–72^.

One could similarly use energy balance models to guide experiments pinpointing the aspects of energy metabolism altered by any diet and leverage untargeted metabolomics to identify putative effector molecules. Beyond detecting microbiome-derived effectors, our methods can be broadly applied to any putative effectors that can be fed to flies as a solid agar. Indeed, this approach is a means to screen drug candidates *in vivo*, many of which interact with the microbiome^56,57^, exemplified by our demonstration that metformin, a drug known to alter human energy metabolism, impacts lactic acid signaling. Furthermore, given the increasing prevalence of metabolic disease in humans^73^, it will be informative to leverage this platform to identify potential protective mechanisms of both microbe-derived and xenobiotic metabolites on energy metabolism in *Drosophila* models of metabolic disease.

## Methods

### Drosophila husbandry

Mated adult female flies 5-7 days post eclosion were used for all experiments. All flies were synchronized on diets during the L1 stage and raised at a density of 50 larvae per 10 mL food at 25 °C on standard cornmeal, molasses, yeast medium. Adult flies were maintained at a density of 10 flies per vial. All flies used for experiments were conventionally reared (not germ free).

### Drosophila strains

We used the following fly strains from the Bloomington *Drosophila* Stock Center: *Oregon-R* (#5), *w*^1118^ (#3605), *Ldh*^16^ (#94698), *tubulin-GAL4* (#5138), *tubulin-GAL80^ts^* (#7017), and *UAS-Ldh-RNAi* (#33640). All relevant genotypes are listed in the figure legends.

### Bacterial culturing

To prepare spent media, *Lp* and *Ap* were grown from frozen stocks into colonies on De Man-Rogosa-Sharpe (MRS) agar plates and individual colonies were grown in liquid MRS medium for 48 h. These cultures were diluted 1:200 in a 50:50 mixture of MRS and CDM and grown for 24 h. For *LpAp* co-cultures, equal volumes of *Lp* and *Ap* MRS cultures were combined and diluted 1:200 in a 50:50 mixture of MRS and CDM. These cultures were then diluted 1:200 in CDM, grown for 24 h, diluted again 1:200, and grown for 48 h. Fresh CDM medium (as a control) and 48 h CDM cultures were pelleted via centrifugation at 4500*g* for 5 min. For 2X media, concentrated CDM was used to culture bacteria. The supernatants were filter sterilized using 0.22 μm filters (Thermo Scientific, 725-2520) to generate CDM control supernatant or cell-free spent media. To create a solid medium for fly ingestion, we mixed CDM supernatant or spent media 1:1 with 2% agar to create a 1% CDM or spent medium agar that was fed to flies. CDM was made according to published protocol^38^. The following bacteria strains were used in this study: *Lp*^40^, *Ap*^31^, *Lp^WCFS1^*^74^, *Lp*^Δ^*^Ldh^* ^75^. All culturing was performed at 30 °C with shaking.

### Colony forming units

To enumerate bacterial density during starvation, flies were initially inoculated with *Lp* and *Ap*. Specifically, flies were allowed to mate for 2 days post eclosion before transfer to media containing a cocktail of antibiotics (neomycin (Fisher Scientific, BP266295), 0.1 mg mL^-1^; ampicillin (Sigma-Aldrich, A9518), 0.1 mg mL^-1^; vancomycin (MP Biomedicals, 02195540.2), 0.1 mg mL^-1^; metronidazole (MP Biomedicals, 0215571025), 0.5 mg mL^-1^) for 3 days. Flies were then transferred to sterile media inoculated with 100 μL of *LpAp* co-culture for 2 days. Flies were then transferred to 1% agar media before collection at various time points post starvation. Flies were surface sterilized using ethanol before homogenization using 0.5 mm silica beads (BioSpec Products, 11079105z) with shaking (Omni International, BeadRuptor 12). Colonies were grown via serial dilution plating of fly homogenate and quantified using the ImageJ plugin Count-On-It^24^.

### Survival assay

Flies were reared as described above and transferred in groups of 10 per biological replicate to experimental media 5-7 days post eclosion. Survival was quantified at 12 h intervals by enumerating dead flies. All experiments were performed at 25 °C, except for temperature-sensitive experiments, which were performed at 29 °C, and vials remained in incubators until all flies were dead. Mean survival was calculated at each time point. The following chemicals were used to feed flies for survival assays: DL-lactic acid (Sigma-Aldrich, L6661), D-glucose (Fisher Scientific, D16-3), agar (Sigma-Aldrich, A1296), and metformin (Sigma-Aldrich, PHR1084). Each biological replicate represents one vial of 10 flies.

### Taste preference

Taste preference was determined using a two-choice assay developed in a previous study^76^. In brief, groups of 10 mated female flies 5-7 days post eclosion were starved for 12 h. Flies were then transferred to vials containing six 10 μL drops of tastant with alternating red (0.5 mg mL^-1^ Amaranth Red, Sigma-Aldrich, A1016) and blue color (0.125 mg mL^-1^ Erioglaucine Blue, Sigma-Aldrich, 861146). Each drop contained a 1X concentration of CDM supernatant or spent media tastant or water. To control for color preference, we alternated each tastant in color across experimental vials to ensure that half of all replicates per group had each tastant in red and half in blue. Flies were fed for 2 h at 25 °C, before freezing at −20 °C. Abdomen color was then manually scored as red, blue, purple, or none using a dissection scope. Preference index was calculated as ((# flies with tastant 1 color)-(# flies with tastant 2 color))/(# flies of any color). Vials with less than 30% of flies scored as any color were not included. For longitudinal measurements of taste preference during starvation, flies were collected at indicated times post starvation and assayed as above.

### Food intake

Food intake was quantified using the <u>ca</u>pillary <u>fe</u>eder assay (CAFÉ) according to previous studies^76,77^. In brief, vials containing 10 mated female flies 5-7 days post eclosion were starved for 4 h. Flies were then transferred to vials whose plugs had two accessible capillary tubes (Drummond Scientific, 2-000-020) containing liquid media as indicated. To more easily visualize liquid volume in capillary tubes, each liquid media contained 0.125 mg mL^-1^ Erioglaucine blue dye. To control for liquid evaporation, an empty vial containing capillary tubes but no flies was included. Evaporation volume was subtracted from each value before analysis. All assays were performed at 25 °C.

### Activity monitor

Activity and sleep were quantified as previously described^78^. Briefly, 5-7 day post-eclosion mated female flies were loaded singly into capillary tubes compatible with the *Drosophila* activity monitor (DAM, Trikinetics)^79^. Each capillary tube (Trikinetics, PPT5×65) contained the indicated media. Activity bouts were collected for 2 days with sleep defined as 5 min of inactivity. Flies with protracted inactivity were determined to have perished and were omitted from analyses. Analysis was performed using the web-based application ShinyDAM^80^.

### Fecundity

The number of eggs laid was enumerated as previously described^81^. In brief, 5-to 7-day old mated female flies were transferred onto indicated media and the number of eggs laid over 24 h was quantified per group of 10 flies.

### Triglyceride quantification

Triglycerides were quantified as previously described^82^. In short, 5-7 day post-eclosion mated female flies were collected at indicated time points in groups of five and homogenized in 200 μL of 0.1% Tween in PBS using 1 mm glass beads (Sigma-Aldrich, Z250473) with shaking (Omni International, BeadRuptor 12). Triglyceride concentration was determined using a colorimetric assay as previously described^83^. Triglyceride concentration was normalized to body weight (Mettler Toledo, MS204S) to calculate percent body fat.

### Mass spectrometry

Liquid chromatography with tandem mass spectrometry was performed as previously described^84^. Briefly, spent media were generated and stored at −80 °C. Samples were split into two 20 μL aliquots and dispensed into 96-well polypropylene plates (Greiner Bio-One, 655161) on ice. To avoid biases associated with repeated freeze-thaw cycles, samples were kept on ice and analysis was performed upon their first thawing. As a quality control step, an additional pooled sample was generated by mixing 5 μL of each sample; this mixed sample was injected at various times throughout LC/MS analysis to check instrument performance. All samples were analyzed in both ionization modes. Sample order was randomly determined for injection to prevent bias in data collection. Blanks consisting of water in place of fresh CDM supernatant or spent media were prepared by performing a parallel extraction and ran as additional quality controls. Chromatographic conditions were modified to shorten the analysis time. Specifically, a shorter ACQUITY BEH Amide 130 Å, 1.7 µm, 2.1 mm x 50 mm (Waters, Milford MA) column was used. Chromatographic gradient was shortened as such: 0-0.5 min 100% B, shifting to 70% B at 1.95 min, 40% B at 2.55 min, returning to 100% B at 3.15 min and held until 3.8 min. All other analytical conditions were as described in Ref. ^84^.

### Metabolomics analysis

Untargeted feature extraction was performed with MS-Dial v. 4.60^85,86^. Mass accuracy tolerances were set to 0.01 Da and 0.015 Da for MS1 and MS2, respectively. Peak height threshold was set to 50k. Lactic acid was annotated based on matching accurate mass, fragmentation spectra, and retention time with an authenticated standard. All reported features were present in 100% of replicates from at least one condition and had a maximum peak height at least five fold greater than the average of the water blanks.

Fold change of a given feature was computed as the ratio of biological replicate means. To prevent division by zero when computing fold changes, we added a pseudocount equal to the lowest nonzero feature area across all samples. For principal component analyses (PCA), we first preprocessed the data by removing features that were constant across all samples of interest and then standardizing all features by subtracting the feature mean and dividing by its standard deviation. PCA was performed using the *PCA* function within scikit-learn using the full exact singular value decomposition solver. Details of statistical testing of normalized peak areas are given in the figure legends.

### Survival curve data analyses

The survival curve *S*(*τ*) is related to the distribution of death times *P*(*τ*) by

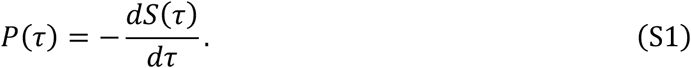

Using this relation, the mean and variance of survival times can be estimated from *S*(*τ*) as

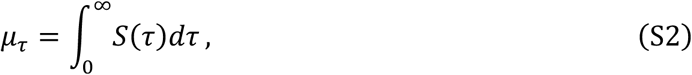

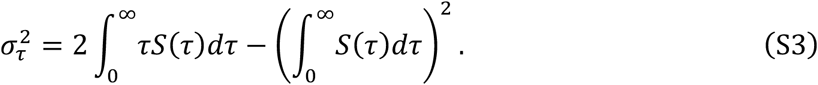

We use Simpson’s method to numerically integrate the empirical survival curves in each vial. In vials in which the flies died very rapidly, we found that the estimator for variance can occasionally result in a slightly negative variance. In such cases, we set the variance to zero. Descriptions of the statistical testing methods used for survival curve moments are given in the figure legends.

### Generalized fly energy balance model

Here, we present a general form of the simplified energy balance model shown in the main text. In this section, we introduce the model and show how it can be reduced to the expressions in the main text. In the following sections, we explore more complex variations of the model.

Drawing upon energy balance models developed for humans^60,87^, we model a population of flies, indexed by *i*, that each have total stored energy *E*_*i*_(*t*) encompassing fat and other energy storage molecules. For simplicity, we treat all energy storage forms as equivalent. We denote the initial stored energy of a fly as *E*_*i*_(*t* = 0) = *E*_*i*,0_. Initially, the population has mean(*E*_*i*,0_) = *μ*_*E*_ and 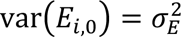. Each fly expends energy at rate *Ø*_O,*i*_(*t*) and intakes energy at rate *Ø*_*I*,*i*_(*t*). The dynamics governing the energy balance of an individual fly are therefore

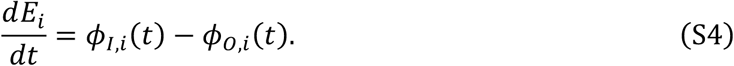

The solution to Eq. S4 is

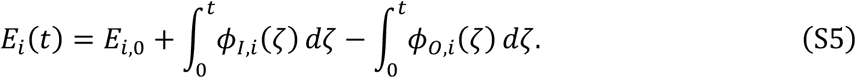

We assume that a fly dies of starvation once its energy storage reaches zero at a starvation time *τ*_*i*_, *E*_*i*_(*τ*_*i*_) = 0.

To reach the minimal null model shown in the main text, we make two simplifications: assuming (1) constant energy intake and expenditure, and (2) the same energy intake and expenditure for all flies in the population, such that *Ø*_*I*,*i*_(*t*) = *Ø*_*I*_ and *Ø*_O,*i*_(*t*) = *Ø*_O_. With these assumptions, the solution is *E*_*i*_(*t*) = *E*_*i*,0_ + (*Ø*_*I*_ − *Ø*_O_)*t* and the starvation time of an individual fly *τ*_*i*_ is

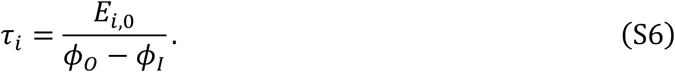

The model predicts that the survival time will be linear as a function of stored energy, a prediction that approximately holds in more complex human energy balance models^87^. Both the mean and variance are based on linear operators (i.e., mean(*αX*) = *α* mean(*X*) and var(*αX*) = *α*^2^ var(*X*)), and thus estimates of the moments of *τ*_*i*_ given *μ*_*E*_ and 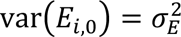 are

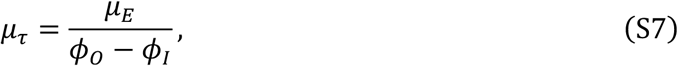

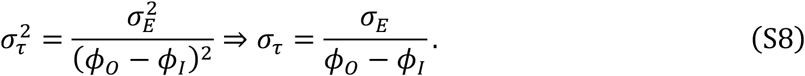

From Eq. S7, we can derive a variable transformation in which there is a linear relationship between inverse mean starvation time and the energy intake *Ø*_*I*_ (and similarly for starvation standard deviation in Eq. S8):

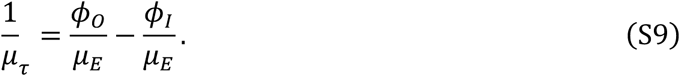

These transformed moments are the most suitable variables for plotting comparisons of the energy balance model to experimental data. We plot a transformed version of Fig. 2C,E in Fig. S8, which shows the agreement between model and experiment.

To relate the above expressions to food energy density *u*, we assume a constant, nutrient-independent inflow of food *Q*, such that *Ø*_*I*_ = *Qu*. In this simplified model, we expect a linear relationship between inverse mean survival time, inverse survival time standard deviation, and food energy density.

This mathematical model indicates that the sensitivity of the starvation assay to changing dietary energy can be superlinear under certain conditions. This superlinearity is apparent in the derivative of the mean survival time with respect to input energy:

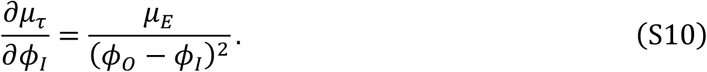

This derivative diverges as the input energy flux approaches *Ø*_O_. Note that the sensitivity will decrease sharply if lifespan plateaus at a finite value.

### Energy balance models with finite lifespan

The model in the previous section predicts that fly lifespan will be infinite once the fly’s energy needs are met. In reality, lifespan will plateau at some finite value due to other factors, limiting the linear relationship between lifespan and food energy density to what we term the “starvation regime”. We introduce two models that produce finite lifespan as examples, but our analyses in this manuscript do not depend on the underlying mechanism for the lifespan plateau.

First, consider a case where the fly’s ability to extract energy from food is limiting (e.g., due to absorption limitations in the gastrointestinal tract): that is, *Ø*_*I*_ ∝ *f*(*u*), where *f*(·) is a saturating function. In this case, both the mean survival time and survival time standard deviation will plateau at some maximum value, mimicking what is seen in experiments.

Another possibility is that lifespan becomes limited by a second factor that is depleted during starvation (e.g., vitamins or nitrogen). The storage level of this compound in fly *i*, which we call *X*_*i*_, should obey similar dynamics to the energy storage:

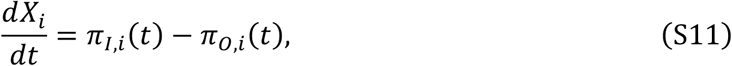

where *π*_*I*_ and *π*_O_represent expenditure and intake, respectively, of the second limiting factor. In this case, lifespan will be dictated by which factor is depleted first. As energy intake is increased, lifespan will smoothly reach a plateau as the other factor becomes limiting. We expect a different variance relationship from the limited energy extraction model as the variance will shift from being controlled by energy storage variance to the storage variance of the second limiting factor.

Note that outside of the starvation regime we consider in this work, the relationship between energy intake and lifespan is not straightforwardly controlled by simple energy balance, with reduced caloric intake associated with longer lifespan in well-fed flies^19^.

### Energy balance model with population-varying energy intake and expenditure

We have thus far assumed that there is only substantial variation in the initial energy storage of the flies. However, variation in energy expenditure and intake is also possible. To determine how such variation would influence survival time, we start from Eq. S5 and consider variation in the net rate of energy expenditure Δ*Ø*_*i*_ = *Ø*_O,*i*_ − *Ø*_*I*,*i*_, with mean(Δ*Ø*_*i*_) = *μ*_Δ*Ø*_ and 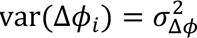. Thus, we aim to determine the value of mean 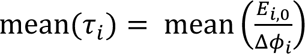. To approximate this quantity, we perform a second-order Taylor expansion for a bivariate function *f*(*X*, *Y*) about the point (*X*, *Y*) = (*α*, *b*):

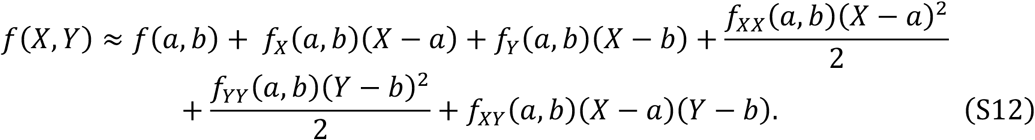

where *f*_*X*_ and *f*_*XX*_denote the first and second derivative, respectively, of *f* with respect to *X*. In our case, *f*(*E*_*i*,0_, Δ*Ø*_*i*_) = *E*_*i*,0_/Δ*Ø*_*i*_ and we expand about *μ*_*E*_ and *μ*_Δ*Ø*_. Substituting in these values and applying the expectation operator to both sides of Eq. S12 yields

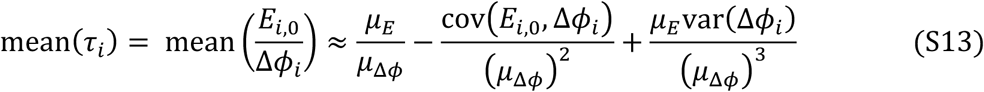

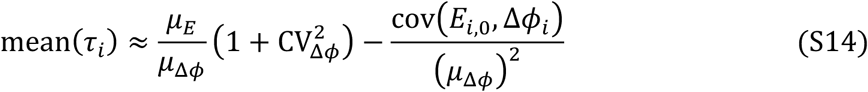

Thus, variation and covariation in Δ*Ø*_*i*_ alter the mean only via higher-order terms. Since *τ*_*i*_ is linearly dependent on *E*_*i*,0_, variation in *E*_*i*,0_ does not alter the mean of *τ*_*i*_.

### Fit of energy balance model to survival data

To fit energy balance models to our experimental data, we used a two-stage approach. First, we fit the non-saturating version of the model to experiments with 0X-2X CDM, as our data indicated that these conditions are outside the lifespan saturation regime. The estimated moments from each vial were treated as independent replicates. We first fit Eq. S7 to the mean survival times. Since the three parameters only appear in ratios, e.g., *μ*_*E*_/*Ø*_O_ and *μ*_*E*_/*Ø*_*I*_, we arbitrarily set *μ*_*E*_ = 1 and estimated *Ø*_O_ and *Ø*_*I*_. We performed this fitting using the MATLAB function *fit*, which minimizes the sum of squared errors. Confidence intervals were estimated using the MATLAB function *confint*. We then fit the variance data using Eq. S9, using estimates of *Ø*_O_ and *Ø*_*I*_ from the mean survival time fit as fixed parameters and *σ*_*E*_ as a fitting parameter.

We found that the simple energy balance model fit the mean survival time data well for flies fed CDM or a pure glucose diet, with adjusted *r*^2^ values of 0.99 and 0.78, respectively. The standard deviation of survival time was also well explained, with adjusted *r*^2^values 0.85 and 0.72. As expected, the naïve energy balance models poorly explained the *Lp*- and *LpAp*^co^-spent media data, with both fits exhibiting negative adjusted *r*^2^ values.

To estimate the parameter changes required to achieve a given change in survival time (e.g., the increase in energy intake *Ø*_*I*_ required to produce the increase in survival between CDM and *Lp*-spent medium), we used Eq. S7 in conjunction with parameters from the above fitting procedure. As an example, we analyze the comparison between 1X CDM versus 1X *Lp*-spent medium. We begin with the parameters estimated using the CDM data: 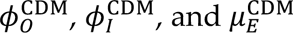. Given that the ratio of mean survival times between these two conditions is 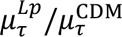, we wish to determine the changes in the energy metabolism parameters (i.e., the values of 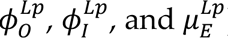) required to produce this change. We analyzed each parameter independently while holding the other two constant. The mean stored energy is

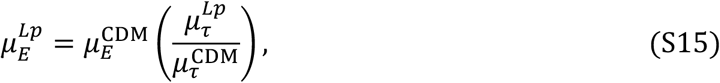

the outgoing energy flux is

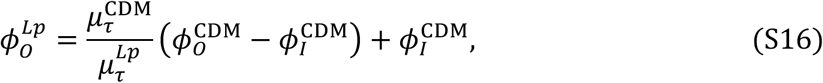

and the incoming energy flux is

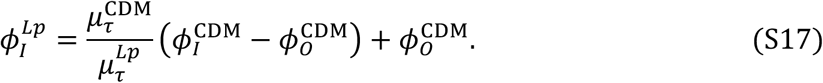

### Linear pharmacokinetic (PK) model of glucose and lactate metabolism in Drosophila

The goal of this analysis is to identify a minimal set of assumptions needed to recapitulate three key experimental observations: (1) a mixture of lactic acid and glucose can promote longer survival than glucose or lactic acid alone, (2) the survival benefit of the mixture saturates with increasing caloric content, and (3) these behaviors are not affected by strong but partial *Ldh* knockdown. Additionally, the model should exhibit regimes in which lactic acid alone promotes similar mean survival as glucose alone. Note that given the vast space of possible models, our goal is to identify whether a model is consistent with our data, not whether it is the only model that can explain the data.

We begin with a two-compartment pharmacokinetic model that considers the gut as one compartment and the rest of the fly body as another compartment. Note that as we are primarily interested in qualitative model behavior, we assume all compartments have the same volume and do not present volumes in our equations. The glycolysis-OXPHOS reaction scheme is simplified and inspired by a previous study^88^. The two primary reactions we consider are:

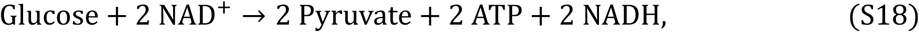

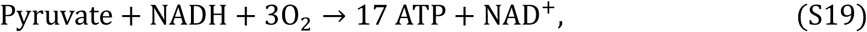

corresponding to the net reactions of glycolysis and OXPHOS, respectively. We also consider the activity of Ldh:

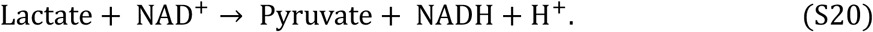

We assume all reactions are linear for analytic tractability. We do not consider the energetic contribution of fat, focusing on food energy extraction, which appears to supply the majority of the fly’s energy in 1X nutrient conditions. Here, we do not explicitly model redox (e.g., NAD^+^/NADH ratio) and assume that the fly is able to maintain this ratio to enable metabolism; we later will consider explicit redox dynamics. We also assume that concentration of ADP, phosphate, oxygen, protons, carbon dioxide, and water do not control the rate of these reactions and thus we do not explicitly model their concentrations.

We model the gut as a well-mixed compartment with constant and equal inflow and outflow rate *δ*. 1/*δ* reflects the passage time of the fly gut. The concentration of glucose and lactic acid in the diet is [G]_diet_ and [L]_diet_, respectively. The concentration in the gut lumen is [G]_gut_ and [L]_gut_, respectively. Absorption of gut glucose and lactic acid occurs linearly at rates *k*_a,G_ and *k*_a,L_, respectively. In the body, lactic acid can be converted to pyruvate by lactate dehydrogenase at rate *k*_Ldh–f_, and pyruvate is converted to lactate at rate *k*_Ldh–r,_. Pyruvate can be used for OXPHOS at rate *k*_OXPHOS_, generating *Y*_OXPHOS_ = 17 ATP per pyruvate. Two pyruvates and *Y*_glycolysis_ = 2 ATP per glucose are produced by glycolysis at rate *k*_glycolysis_. Glucose, lactate, and pyruvate are cleared from the body or used for other metabolic reactions at clearances CL_σ_. As compartment volume is implicit in our model, the clearance processes are represented as first-order elimination constants *k*_e,σ_ = CL_σ_/*V*. We assume the incoming energy flux *Ø*_*I*_ is proportional to the ATP generated by glycolysis and OXPHOS. The dynamics are thus

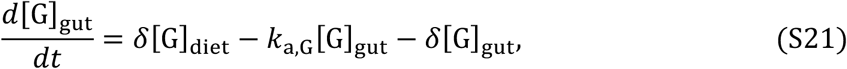

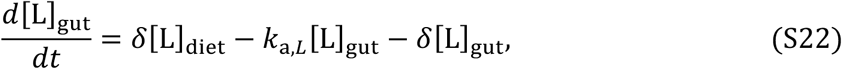

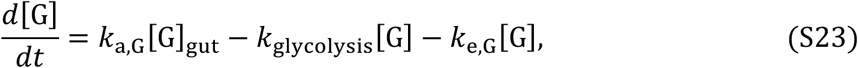

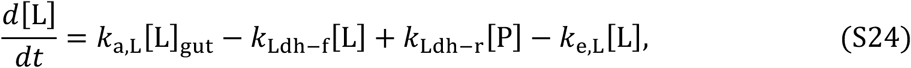

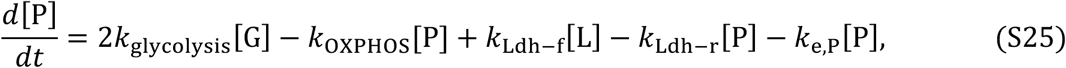

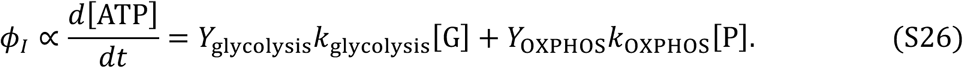

### Steady-state behavior of the linear lactic acid and glucose PK model

We sought to determine whether the above linear model could reproduce the starvation dynamics seen in our assays. Assuming that the fly’s internal reaction network reaches steady state on a timescale faster than starvation, the energy influx from diet will be

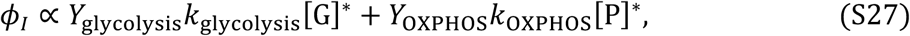

where [G]^∗^ and [P]^∗^ denote the steady-state concentrations of glucose and pyruvate, respectively, in the central compartment. Since this model is linear, we can solve for these quantities by setting the left-hand-side of Eq. S21-S26 to zero:

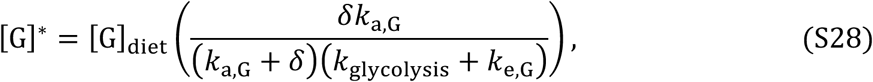

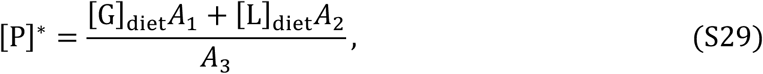

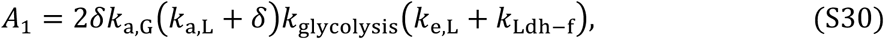

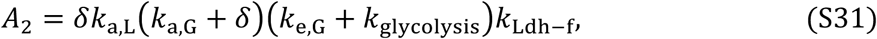

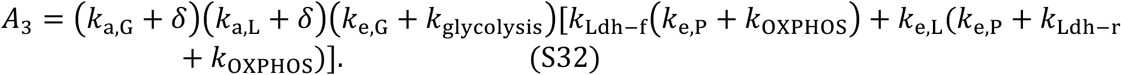

Note that the energy flux is a linear function of the input dietary compositions [G]_diet_ and [L]_diet_. In our experiments, we impose a linear relationship between these two compositions to maintain approximately equal caloric density (Δ[G]_diet_ = −2Δ[L]_diet_). Thus, this model is not capable of reproducing the nonlinear trends we see in our assay in which a mixture of glucose and lactic acid prolongs lifespan more than diets of either compound alone.

### Genome-scale metabolic model of glucose and lactic acid metabolism

The linear model analyzed above is a substantial simplification of *Drosophila* metabolism, which involves thousands of reactions and factors such as redox balance. Before we turned our attention to models with active regulation of metabolic activity, we first sought to determine whether a more realistic model of metabolism could explain our experimental results. To do this, we analyzed iDrosophila1, a genome-scale metabolic model (GEM) of *Drosophila* metabolism^45^. The goal of a GEM is to represent the complete set of metabolic reactions that can occur within an organism, typically constructed by referencing the organism’s annotated genome. GEMs can then be used to model metabolism using flux-balance analysis (FBA), which predicts the metabolic fluxes in an organism given an objective function such as biomass production. There are significant differences between the GEM model formalism and the PK model formalism. While the PK model specifies a set of reaction kinetics and estimates the steady-state chemical and energy fluxes given a diet, FBA-based modeling finds a flux configuration maximizing an objective function, ignoring reaction kinetics. Another notable difference is that iDrosophila1 models whole-organism growth, considering factors such as amino acids that are not explicitly accounted for in our energy-based PK model. However, this reliance on non-energetic factors is represented in the version of the energy balance model with a second limiting factor discussed in a previous section. Despite these differences, iDrosophila1 still provides a useful benchmark for our simple PK model. If our basic assumptions about energy metabolism are correct, the two models should produce similar behavior despite the vast differences in complexity.

As a baseline diet for the iDrosophila1 analysis, we used metabolite intake values calculated for the Holidic defined diet^39,89^. The primary carbon source in this diet is sucrose, a disaccharide composed of glucose and fructose. To maintain a similar dietary energy composition, we replaced one unit of sucrose flux with two units of glucose flux. Similarly, one unit of sucrose flux was replaced with four units of lactic acid flux. We altered diet composition by changing the maximum uptake rates of these carbon sources. Note that while our experiments indicated that that lactic acid and lactate are not equivalent, with only lactic acid extending starvation survival, iDrosophila1 only considers L-lactate. L-lactate supports similar growth as other carbon sources in iDrosophila1.

In Fig. S9, we show the predicted growth rates from iDrosophila1 supplied the Holidic diet with mixtures of glucose and L-lactate as the primary carbon sources. We used the *gurobi* solver in the MATLAB version of the COBRA toolbox to solve for the optimal flux configuration and growth rate. The “1X” condition corresponds to the maximum carbon flux in the original Holidic diet, while “2X” corresponds to double that flux. iDrosophila1 predicted similar behavior as the linear PK model, with a linear relationship between glucose fraction and growth. Similar to the PK model, glucose resulted in a slight growth advantage over L-lactate. Consistent with our energy balance models, iDrosophila1 predicted saturation of growth rate as overall carbon flux was increased: the 2X carbon condition did not provide double the growth rate in the 1X condition. This saturation occurs as growth under the Holidic diet is limited by essential amino acids, not carbon or energy influx^89^.

From this analysis, we conclude that our experimental results cannot be straightforwardly explained by a more realistic model of the *Drosophila* metabolic network. Despite only encoding a few reactions, our linear PK model performs similarly to a GEM encoding 8,230 reactions.

### Pharmacokinetic-pharmacodynamic model of Drosophila energy metabolism incorporating OXPHOS regulation

Our analysis of the linear PK model and the iDrosophila model strongly indicates that some form of diet-dependent regulation is required to produce the trends seen in our experiment. Thus, we explored a pharmacokinetic-pharmacodynamic model in which dietary components can alter the regulation of energy metabolism. We considered a Gaussian function in which the ratio of glucose and lactic acid within the gut compartment modulates OXPHOS activity:

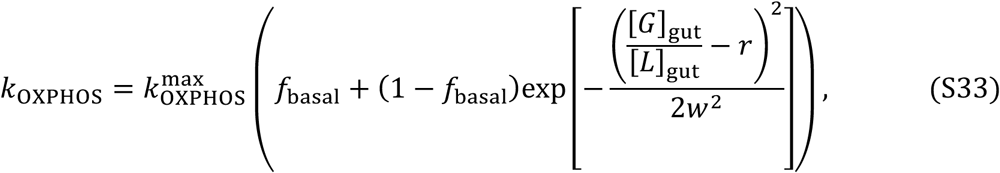

where 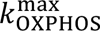 is the maximum OXPHOS rate, *f*_basal_ is the fraction of OXPHOS activity that is basally active, *r* is the ratio of gut glucose to lactic acid at which OXPHOS activity is maximized, and *w* is a parameter controlling the width of the regulation function. This function produces a bell curve response, with OXPHOS activity maximized at 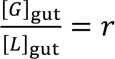 and minimized when only glucose or lactic acid are supplied.

Given that our goal is a qualitative recapitulation of the phenomena seen in our experimental starvation assays, we did not attempt to quantitatively parameterize the model based on our experiments. Owing to the large number of parameters in this model, such parameterization likely would require an intractable number of additional experiments. Instead, to analyze the model, we numerically integrated the ODEs until all variables reached a steady state using MATLAB’s *ode15s* integrator. We found that this model can qualitatively recapitulate the three key phenomena observed in our starvation assays (Fig. 4E). In the simulations shown in the main text, we assumed that nutrient absorption is rapid relative to gut dilution and that Ldh operates primarily in the forward direction (i.e., generating pyruvate). Table S1 contains a detailed list of all parameters. For the model representing Ldh knockdown, we reduced the value of both the forward and reverse Ldh reaction coefficients by the specified knockdown fraction.

Note that multiple regulatory schemes could produce similar behavior to this model. For example, OXPHOS regulation instead being dependent on the internal compartment glucose ratio [*G*]/[*L*] would produce similar behavior. Non-monotonic regulation in response to only lactic acid levels could also produce similar effects. It is also possible that regulation of other enzymes within the pathway could produce non-monotonicity in survival. For example, in Eq. S28-S32 the portion of energy flux from OXPHOS is dependent on the forward and reverse rate constants of Ldh. Thus, a regulatory function modulating these two rates may exist that can reproduce the non-monotonicity in survival. However, such a regulatory scheme is less parsimonious than the proposed OXPHOS regulatory scheme and inconsistent with prior *in vitro* work finding that glucose and lactic acid mixtures regulate OXPHOS^54^. A scheme in which Ldh is the key regulatory target would also likely be inconsistent with the results of our Ldh knockdown and metformin experiments.

The above PKPD model, while it captures the survival dynamics of flies fed glucose and lactic acid diets, cannot capture the results of the priming experiment as it assumes that regulation is instantaneous. To capture the observed priming results, one could incorporate concepts from indirect response pharmacodynamic models, in which an effector modulates an intermediate factor that then impacts the phenotype of interest^90^.

### Analysis of fat breakdown data

We analyzed the fat breakdown time series using multiple fitting approaches. We first estimated the time to 50% fat breakdown by fitting a piecewise linear model to the time series. We identified the time interval in which the mean remaining body fat percentage crossed 50%. In all time series, this crossing occurred only once. Once we identified this interval, we fit a linear model *y* = *β*_1_*t* + _0_, where *t* is time and *y* is the percentage of remaining body fat, using MATLAB’s *fit* function and computed parameter confidence intervals using the *confint* function. From these parameters, we estimated the time at which the body fat decreased to 50% of its initial value as *t*_50_ = (50 − *β*_0_)/*β*_1_. To estimate the confidence interval of this depletion time, we used the standard error propagation formula: 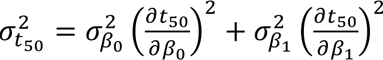.

To obtain the average rate of fat depletion, we fit a linear model to the entire time series using the methods specified above. We fixed the intercept to *β*_0_ = 100 and estimate the depletion rate *β*_0_. From this linear fit, we computed an additional estimate of the *t*_50_.

For estimating the confidence interval of this estimate of *t*_50_, we set 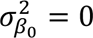. We also fit an exponential model, *y* = *β*_0_*t*^−*β*+*t*^, where again *β*_0_ = 100, using the same MATLAB functions as with the linear model. The exponential model exhibited a somewhat lower adjusted *r*^2^ than the linear model.

While the results varied slightly across these three analysis methods, all supported the same qualitative conclusion: starved flies exhibited the fastest fat breakdown, flies fed pure glucose or lactic acid exhibited an intermediate rate, and flies fed a mixture of glucose and lactic acid exhibited the slowest fat breakdown.

### Modeling acid/base equilibria in fly diets

Our experiments indicated a substantial difference between the effects of lactate/acetate versus lactic acid/acetic acid, with sodium lactate/acetate providing minimal lifespan extension. To understand the composition of diets initially containing the weak acid form of these compounds, we use standard acid/base modeling approaches. Consider a mixture of two weak acids, HA and HB, with logarithmic dissociation constants p*K*_*α*,A_ and p*K*_*α*,B_and initial concentrations [HA]_0_and [HB]_0_, respectively. At equilibrium, a fraction of these acids will dissociate into their conjugate base with concentrations [A^−^] and [B^−^]. Neglecting the small initial concentration of protons present in water, the final proton concentration will be [H^+^] = [A^−^] + [B^−^]. The remaining concentration of weak acid can be found by mass balance given the conjugate base concentrations. At equilibrium, the concentrations will be

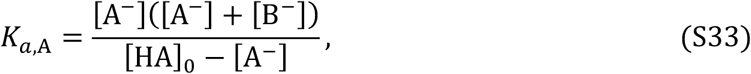

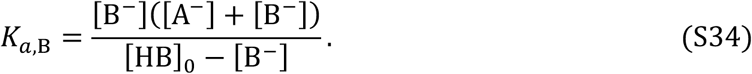

For a given mixture of lactic acid (p*K*_*α*_ = 3.86) and acetic acid (p*K*_*α*_ = 4.76), we numerically solved these equations for [A^−^] and [B^−^] for a range of dietary compositions using MATLAB’s *fsolve* function (Fig. S5C, D). For both single acid and mixed acid diets, the weak acid form of the compound is predicted to dominate at equilibrium.

## Supplementary Text

### Estimation of the maximal caloric content of CDM and lactic acid/glucose diets

Here, we outline our approach for estimating the maximum available energy densities of the diets used in our experiments. We compute the Gibbs free energy of combustion (Δ*G*) of dietary components using eQuilibrator^91^, a calculator for thermodynamic analysis of biochemical reaction systems. This Gibbs free energy represents the maximum possible energy yield from complete oxidation of the dietary component. We calculated these free energies at pH 7, pMg 3, ionic strength 0.25 M, and reactant concentrations of 1 mM. The dominant source of energy and carbon in CDM is glucose, which has Δ*G* = −2911 kJ/mol. Lactic acid fermentation converts a molecule of glucose into two molecules of lactic acid, which each have Δ*G* = −1347 kJ/mol. Thus, complete fermentation of all glucose in CDM would result in only a ∼7% loss in energy. As the spent media are filter sterilized before being fed to flies, we neglected the energy that the fly could gain from digestion of bacterial cells. This 7% difference in energy cannot explain the survival effects of *Lp*-spent medium nor the difference in survival between the diets composed of lactic acid or glucose alone. To mimic the outcome of lactic acid fermentation, we constructed our lactic acid and glucose diets by exchanging glucose with lactic acid at a 1:2 ratio. We refer to the resulting diets as “near isocaloric” since the differences are small relative to the nonlinear glucose-lactic acid interaction we observe.

Note that the increased energy yield we find in the *Lp*-spent medium condition indicates that the actual energy yield of *Drosophila* from 1X CDM diets is substantially lower than the maximum estimated here. Indeed, our model estimates that in the 1X CDM condition, at least ∼45% of the theoretically available dietary energy is left unutilized.

### Estimation of in situ Drosophila microbiome metabolic fluxes

First, we determined a relationship between *Lp* growth and lactate production. A prior study found that in MRS medium *Lp* grows to a density of ∼2 × 10^9^ CFU/mL and produces ∼200 mM of D+L-lactate (Fig. 2A and C)^22^. These values lead to lactate yield estimate of *Y* = 10^−13^ mol/CFU. As lactic acid fermentation is a central metabolic reaction, we assume the yield does not vary substantially between growth conditions.

Next, we estimated the typical population size and growth rate of *Lp* within non-starved *Drosophila.* In our experiments, we found that the steady-state *Lp* population, *ρ*_LP_, within well-fed flies was ∼5 × 10^5^ CFU/óly (Fig. S1). Assuming that the dominant source of death within the fly gut is dilution out of the gut, the steady-state growth rate of *Lp* must match the rate of dilution. The transit time of a fly gut is ∼1 h (Fig. 7D, Ref. ^92^), corresponding to a dilution rate of *δ*∼24 day^−1^. Thus, the estimated per-fly lactate flux is *ρ*_LP_*Yδ* = 1.2 × 10^−6^ mol/day, corresponding to a carbon atom flux of 3.6 × 10^−6^ mol/day.

Finally, we determined whether the above flux is significant relative to the incoming flux of carbon compounds from a typical diet. Our CAFÉ assay data indicated that flies consume ∼1.5 *μ*L/day of food across diets. Well-fed flies consume a diet in which the primary carbon sources are yellow cornmeal (61.3 g/L) and molasses (75.2 mL/L). Yellow cornmeal is 77% carbohydrates by mass (FoodData Central ID: 169697, USDA, 2025).

Molasses is 74.7% carbohydrates by mass and has a specific gravity of 1.33 (FoodData Central ID: 168820, USDA, 2025). The mass density of molasses in the fly food is therefore 101.5 g/L. Thus, an estimate of the total carbohydrate mass density of the food is 122 g/L. The carbohydrates in these two carbon sources are mostly composed of simple sugars or polymers of simple sugars. Assuming all sugars have a composition similar to glucose, this diet contains a glucose-equivalent concentration of 677 mM, corresponding to an input glucose flux of 1 × 10^−6^ mol/day and a carbon atom flux of 6 × 10^−6^ mol/day. The estimated *Lp*-derived lactic acid carbon atom flux of 3.6 × 10^−6^ mol/day is on the same order of magnitude as the dietary carbon flux, indicating that *Lp* cells can potentially produce physiologically relevant fluxes of lactic acid *in situ*.

## Acknowledgements

We thank Dr. François Leulier for sharing the *Lp^WCFS1^* and *Lp*^Δ^*^Ldh^* strains with kind permission from Dr. Pascal Hols. We thank Dr. John Vaughen and Dr. Thomas Clandinin for permitting use of the DAM and help with analyzing data. We are grateful to M. Cesur, K. Patil, R. Porter, B. Fu, and J. Sonnenburg and members of the O’Brien, Huang, Good, and Sonnenburg labs for helpful discussions. We are thankful for Dr. L. Wat’s kind sharing of a cartoon fly illustration. Stocks obtained from the Bloomington Drosophila Stock Center (NIH P40OD018537) were used in this study. We thank the TRiP at Harvard Medical School (NIH/NIGMS R01-GM084947) for providing transgenic RNAi fly stocks used in this study. We acknowledge critical resources and information provided by FlyBase^93^. FlyBase is supported by a grant from the National Human Genome Research Institute at the U.S. National Institutes of Health (U41 HG000739) and by the British Medical Research Council (MR/N030117/1). The authors acknowledge support from a Helen Hay Whitney Postdoctoral Fellowship (to J.W.M.), a Stanford PRISM Baker Fellowship (to J.A.L.), the Bio-X Summer Research Experience for Undergraduates program (to A.M.S.), NSF Awards IOS 2032985 and CAREER 2144342 (to W.B.L.), CZI Grant DAF2020-217689 (to L.E.O. and K.C.H.), NIH Awards R35GM141885 (to L.E.O.) and R35GM146949 (to B.H.G.), and NSF Award EF-2125383 (to. K.C.H.). B.H.G., L.E.O., and K.C.H. are Chan-Zuckerberg Biohub Investigators.

## Data availability

All codes and processed data used to generate figures are available at https://github.com/jamie-alc-lopez/fly_energy_metabolism.

## Supplemental Figures

**Figure S1:**
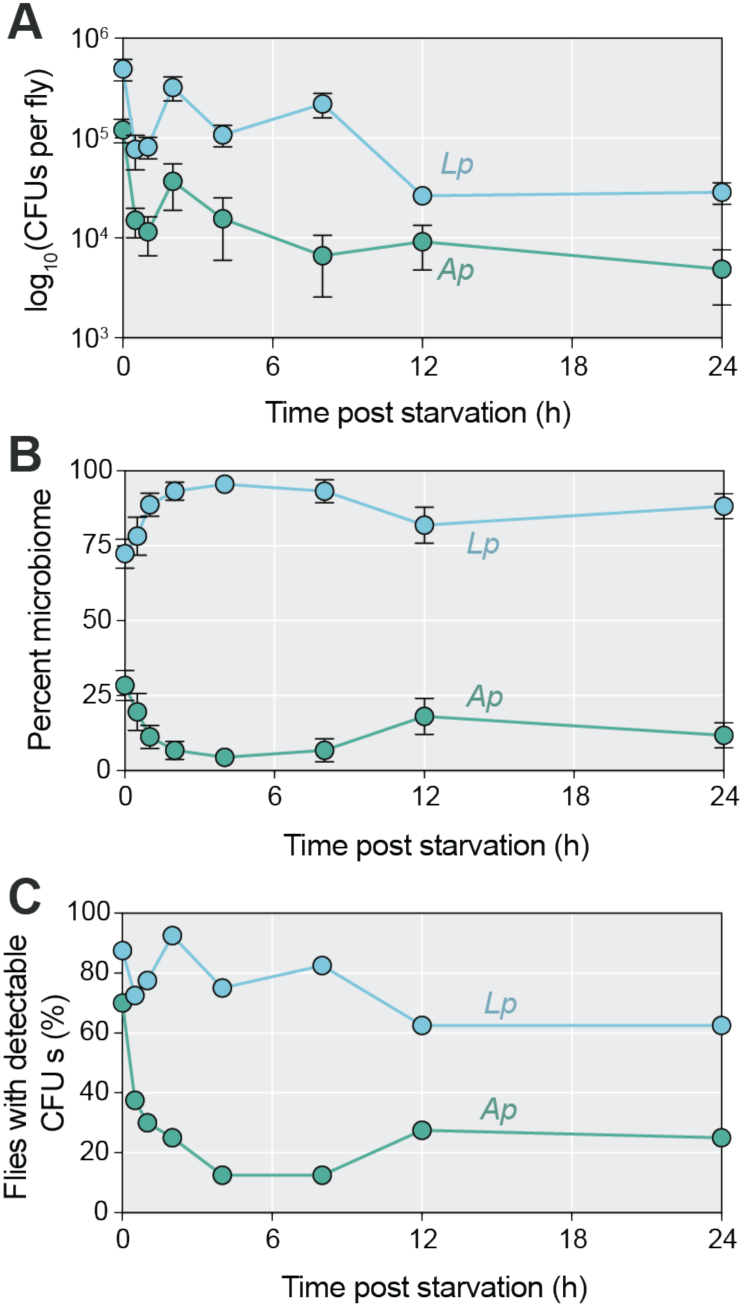
Microbial load decreases during starvation and composition shifts to increased frequency of the *Lactiplantibacillus* genus. A) Bacterial load in the *Drosophila* gut as measured by colony forming units on agar selective for *Lactiplantibacillus plantarum* (*Lp*) or *Acetobacter pasteurianus* (*Ap*) growth over 24 h of starvation. Microbe load decreased for both species. B) Microbiome composition measured as the relative abundance of *Lp* and *Ap* shifts toward increased *Lp* frequency. C) Percentage of flies with any culturable microbes of either species decreased during starvation. More *Lp* cells were culturable at 24 h of starvation than *Ap* cells. *N* = 40 biological replicates.

**Figure S2:**
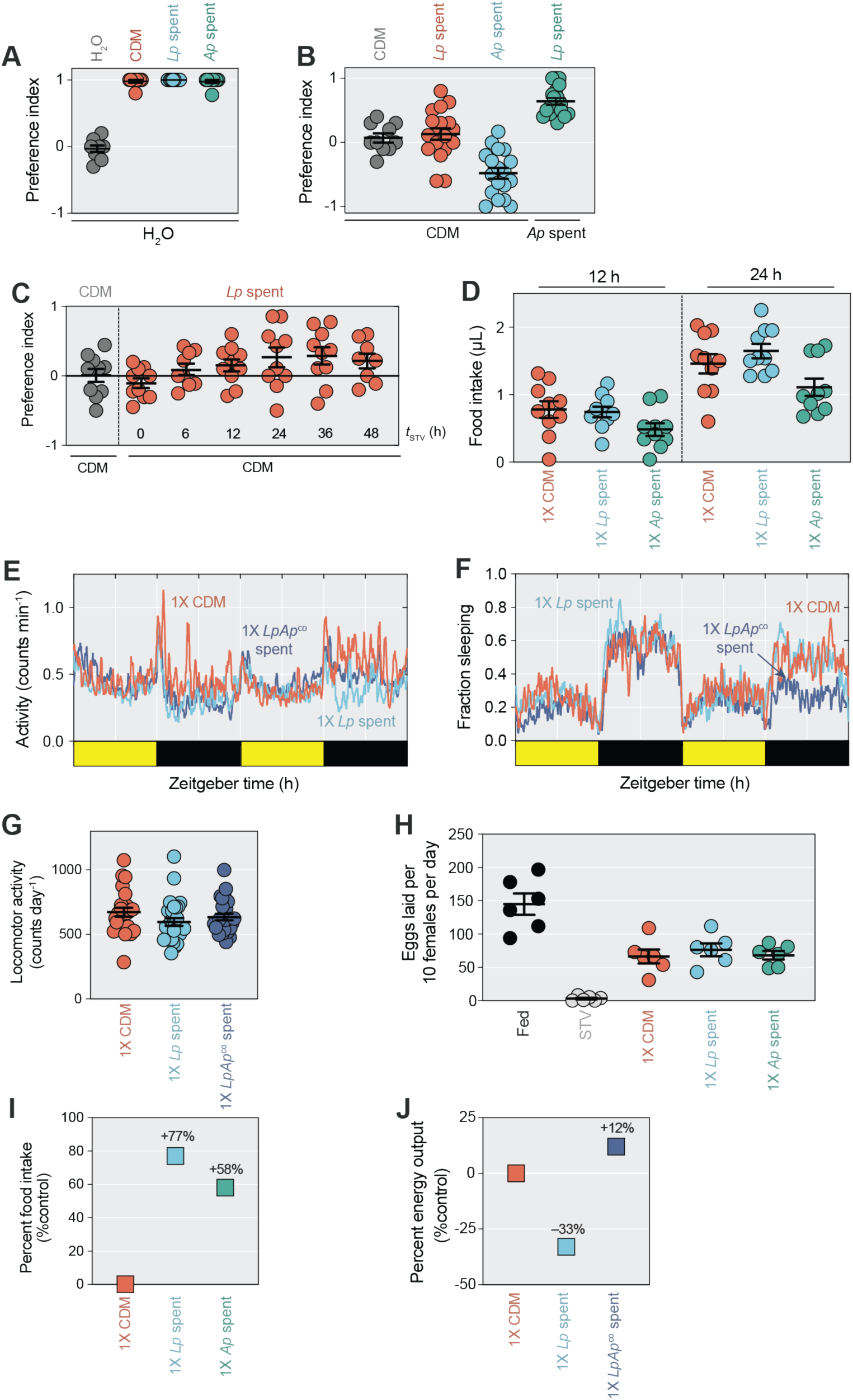
Microbe-derived metabolites do not impact starvation survival via differential food intake, active energy expenditure, or fecundity. A) Taste preference in *Ore-R* mated females as quantified by a dye-based, two-choice feeding assay demonstrated that fresh CDM supernatant, *Lp-*spent medium, and *Ap-*spent medium are appetitive to flies. *N* = 10 biological replicates, *n* = 100 flies per condition. B) Taste preference in *Ore-R* mated females between CDM supernatant and *Lp-* spent medium was neutral, however flies preferred CDM supernatant or *Lp-* spent medium compared to *Ap-*spent medium. *N* = 10-18 biological replicates, *n* = 100-180 flies per condition. C) Taste preference in *Ore-R* mated females between fresh CDM supernatant and *Lp-*spent medium remained neutral during prolonged starvation. *N* = 8-10 biological replicates, *n* = 80-100 flies per condition. D) Food intake did not change in *Ore-R* mated females between flies fed fresh CDM supernatant, *Lp-*spent medium, and *Ap-*spent medium. E) Locomotor activity in *Ore-R* mated females as measured using the *Drosophila* activity monitor (DAM) over 48 h of feeding with fresh CDM supernatant, *Lp-* spent medium, and *LpAp*^co^*-*spent medium. F) Time spent sleeping in *Ore-R* mated females measured using DAM over 48 h of feeding with fresh CDM supernatant, *Lp-*spent medium, and *LpAp*^co^*-*spent medium. G) Daily activity was similar among *Ore-R* mated females fed fresh CDM supernatant, *Lp-*spent medium, and *LpAp*^co^-spent medium. *N* = 24-28 biological replicates. H) The number of eggs laid in 24 h by *Ore-R* mated females fed with CDM supernatant, *Lp-*spent medium, and *Ap-*spent medium was similar. *N* = 6 biological replicates, *n* = 60 flies per condition. I) Energy balance model prediction for the change in food intake necessary to account for the observed survival through a change in energy intake. J) Energy balance model prediction for the change in activity necessary to account for the observed survival through a change in energy output.

**Figure S3:**
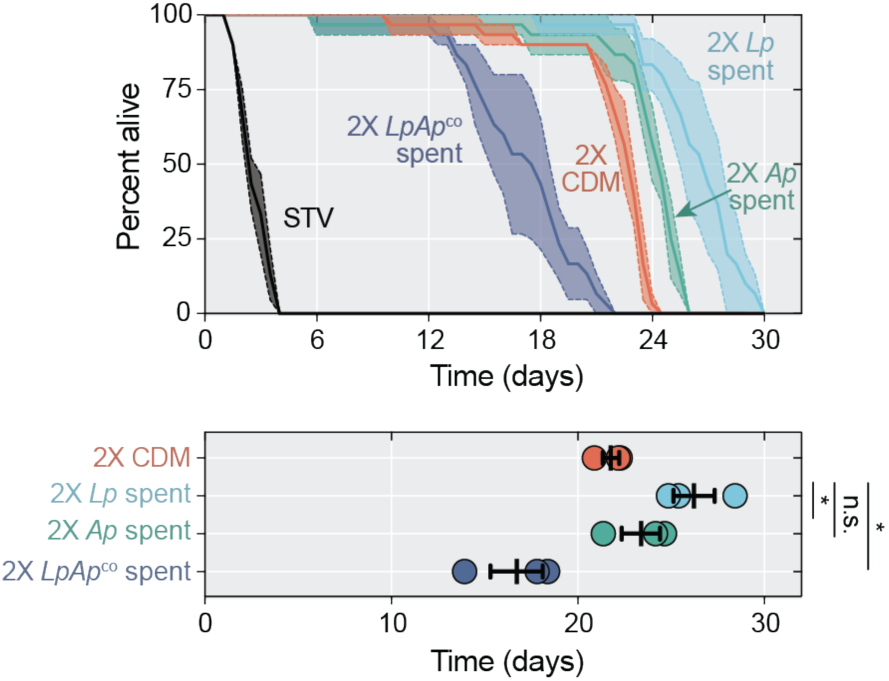
Caloric value affects how spent media impact starvation survival. Mean survival was significantly higher in *Ore-R* mated females fed *Lp-*spent medium compared to CDM supernatant (*p* = 0.043; one-way ANOVA followed by Dunnett’s multiple comparison test). Mean survival was not significantly different between *Ore-R* mated females fed *Ap-*spent medium compared to CDM (*p* = 0.59; one-way ANOVA followed by Dunnett’s multiple comparison test). Mean survival was significantly shorter in *Ore-R* mated females fed *LpAp*^co^-spent medium compared to CDM (*p* = 0.024; one-way ANOVA followed by Dunnett’s multiple comparison test). *N* = 3 biological replicates, *n* = 30 flies per condition. *: *p* < 0.05; ns: not significant; error bars indicate 1 SEM.

**Figure S4:**
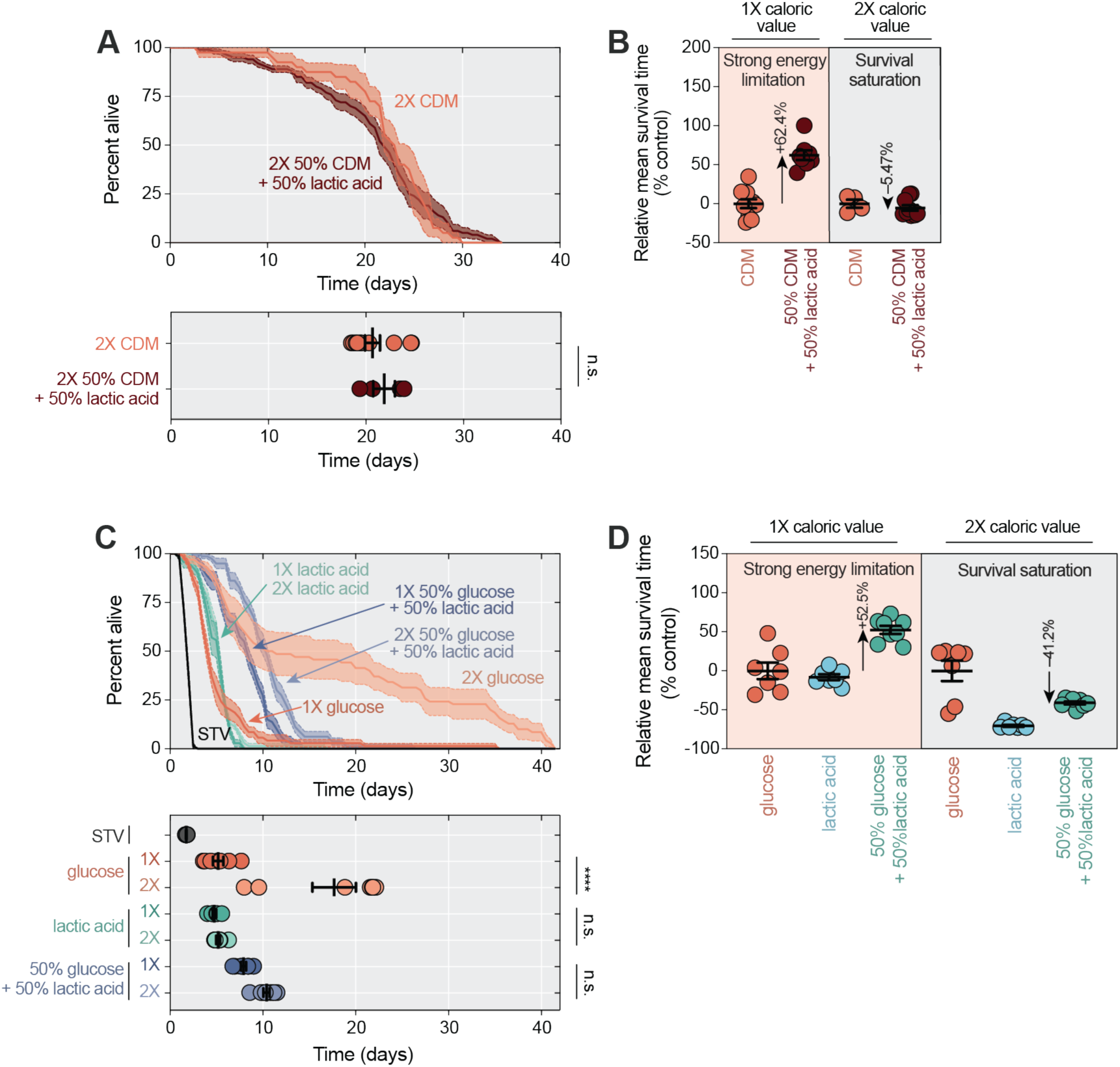
Caloric value affects how lactic acid impacts fly survival. A) Mean survival was not significantly different in *Ore-R* mated females fed 2X 50% CDM + 50% lactic acid compared to flies fed 2X CDM (*p* = 0.41; Student’s *t* test). *N* = 4-10 biological replicates, *n* = 40-100 flies per condition. B) The percent difference in mean survival relative to the control was higher at lower caloric values in the strong energy limitation regime compared with high caloric values in the survival saturation regime for flies fed CDM versus flies fed CDM supplemented with lactic acid. C) Mean survival was significantly higher in *Ore-R* mated females fed 2X glucose compared to flies fed 1X glucose (*p* < 0.0001; two-way ANOVA followed by Sidak’s multiple comparison test). Mean survival was not significantly different between *Ore-R* mated females fed 2X lactic acid compared to flies fed 1X lactic acid or between flies fed 2X 50% glucose + 50% lactic acid compared to flies fed 1X 50% glucose + 50% lactic acid (*p* = 0.99 and 0.18, respectively; two-way ANOVA followed by Sidak’s multiple comparison test). *N* = 7-8 biological replicates, *n* = 70-80 flies per condition. D) The percent difference in mean survival relative to the control was higher at lower caloric values in the strong energy limitation regime compared with high caloric values in the survival saturation regime for flies fed glucose + lactic acid versus control diet flies. ****: *p* < 0.0001; ns: not significant; error bars indicate 1 SEM.

**Figure S5:**
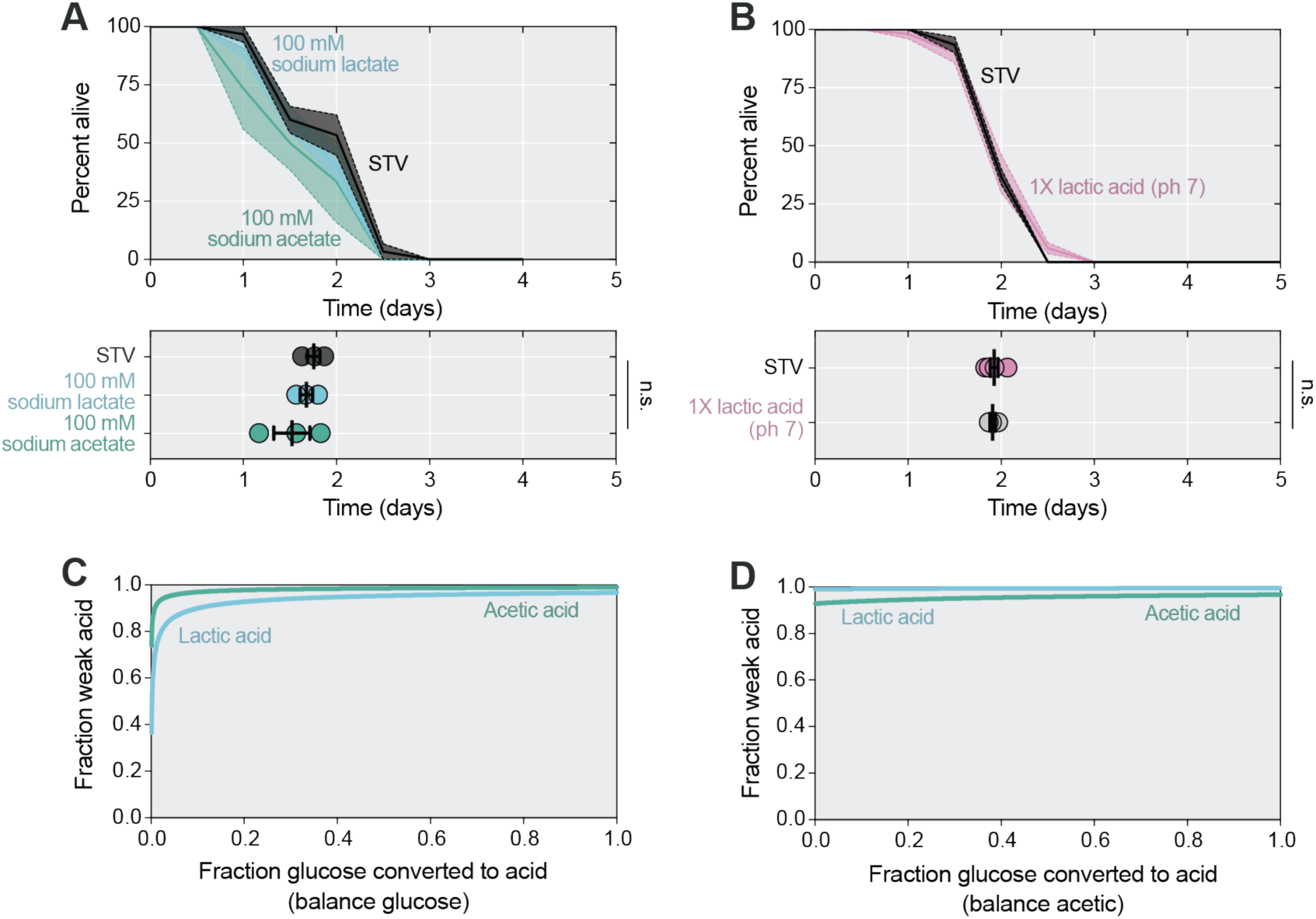
Neither dietary lactate nor neutralized lactic acid extend fly survival. A) Mean survival was not significantly different in *Ore-R* mated females fed 100 mM sodium lactate or 100 mM sodium acetate compared to completely starved flies (*p* = 0.87 and 0.37, respectively; one-way ANOVA followed by Dunnett’s multiple comparison test). *N* = 3 biological replicates, *n* = 30 flies per condition. B) Mean survival was not significantly different in *Ore-R* mated females fed 1X lactic acid corrected to pH 7 with dropwise sodium hydroxide compared to completely starved flies (*p* = 0.80; Student’s *t* test). *N* = 3-5 biological replicates, *n* = 30-50 flies per condition. C) Fraction of lactic acid and acetic acid predicted to exist in the weak acid form across varying acid to glucose ratios in single acid solutions. D) Fraction of lactic acid and acetic acid predicted to exist in the weak acid form across varying acid to glucose ratios in mixed acid solutions.

**Figure S6:**
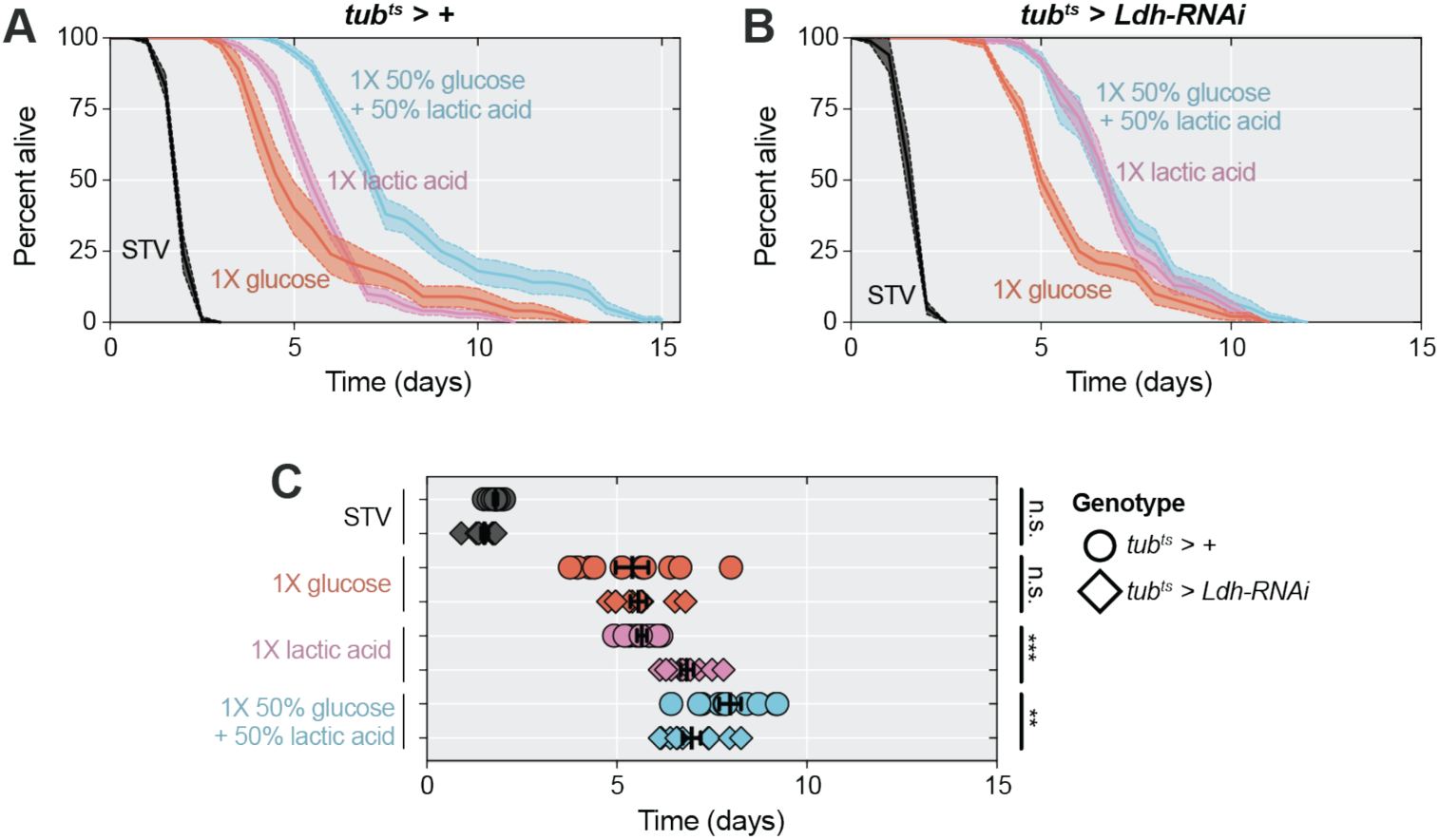
Whole-body Ldh is dispensable for lactic acid-mediated effects on survival. A) Mean survival was significantly higher in *tub^ts^ > +* mated females (genotype: *w^1118^*;; *tub-GAL4, tub-GAL80^ts^*/+) fed 1X 50% glucose + 50% lactic acid compared to flies fed 1X glucose or 1X lactic acid (*p* < 0.0001 for both comparisons; two-way ANOVA followed by Tukey HSD test). Mean survival was not significantly different between *tub^ts^ > +* mated females fed 1X glucose compared to flies fed 1X lactic acid (*p* = 0.86; two-way ANOVA followed by Tukey HSD test). B) Mean survival was significantly higher in *tub^ts^ > Ldh-RNAi* mated females (genotype: *w^1118^*;; *tub-GAL4, tub-GAL80^ts^*/*UAS-Ldh-RNAi*) fed 1X 50% glucose + 50% lactic acid compared to 1X glucose but not 1X lactic acid fed flies (*p* = 0.0003 and 0.98, respectively; two-way ANOVA followed by Tukey HSD test). C) Mean survival was not significantly different between *tub^ts^ > +* and *tub^ts^ > Ldh-RNAi* mated females in either starved or 1X glucose feeding conditions (*p* = 0.39 and 0.61, respectively; two-way ANOVA followed by Tukey HSD test). Mean survival was significantly increased in *tub^ts^ > Ldh-RNAi* compared to *tub^ts^ > +* mated females fed 1X lactic acid (*p* = 0.0005; two-way ANOVA followed by Tukey HSD test). Mean survival was significantly shorter in *tub^ts^ > Ldh-RNAi* compared to *tub^ts^ > +* mated females fed 1X 50% glucose + 50% lactic acid (*p* = 0.0027; two-way ANOVA followed by Tukey HSD test). *N* = 10 biological replicates, *n* = 100 flies per condition. **: *p* < 0.01, ***: *p* < 0.001, ****: *p* < 0.0001; ns: not significant; error bars indicate 1 SEM.

**Figure S7:**
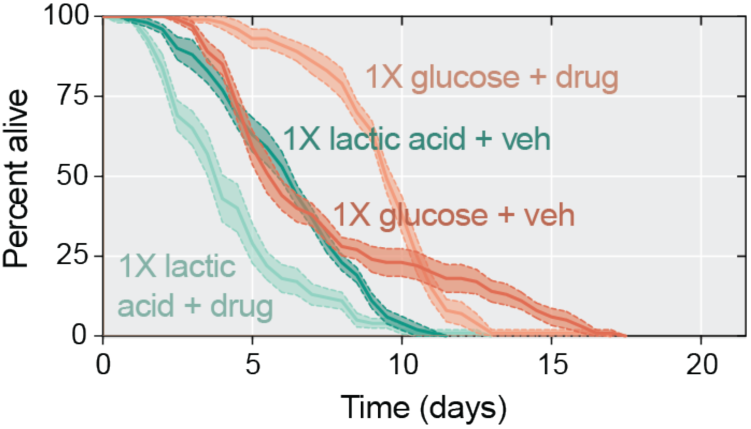
Metformin impacts survival in flies fed glucose and lactic acid. Survival was significantly longer in *Ore-R* mated females treated with 25 mM metformin and fed 1X glucose compared to vehicle-treated flies fed 1X glucose, potentially due to a titrating effect on energy yield from OXPHOS. Survival was significantly reduced in *Ore-R* mated females treated with 25 mM metformin and fed 1X lactic acid compared to vehicle-treated flies fed 1X lactic acid. *N* = 10 biological replicates, *n* = 100 flies per condition.

**Figure S8:**
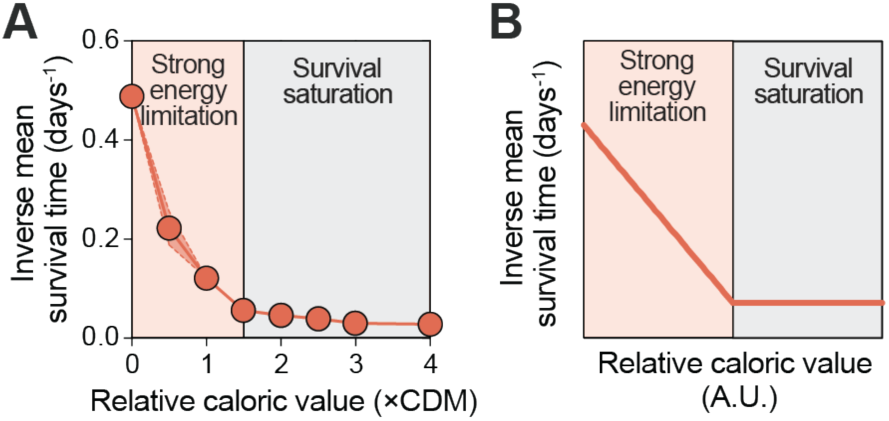
Transformed survival moments of CDM dilution series compared to energy balance model. A) Inverse mean survival is predicted to scale linearly between 0-1.5X CDM and to saturate between 2-4X CDM. B) Energy balance model predictions for inverse mean survival as a function of increasing relative caloric value. Inverse mean survival is predicted to scale linearly in the energy limitation regime and to saturate when energy demands for normal lifespan are met.

**Figure S9:**
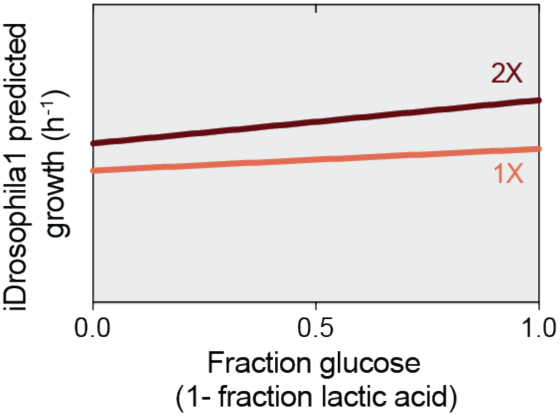
Genome-scale metabolic model of *Drosophila* metabolism (iDrosophila1) predicts a linear relationship between diet composition and overall metabolic flux. Predicted growth rate, a normalized representation of metabolic flux of ATP and other essential compounds, in the iDrosophila1 model supplied with mixtures of only glucose and L-lactate. Diet composition was modified by changing the maximum allowable input flux of these two carbon sources. Aside from input fluxes of sucrose (set to zero), glucose, and L-lactate, the fly diet composition is identical to the Holidic diet condition from previous studies^89^.

**Table S1:** Parameters used in the PKPD model simulations shown in Fig. 4D-F. All rate parameters have units of inverse time. These parameters are not a quantitative fit to the reaction kinetics of metabolism in *Drosophila*, but rather represent a set of parameters that provide a qualitative recapitulation of our data. The linear PK model simulated with these parameters produced qualitatively similar trends to those predicted by the iDrosophila1 genome-scale metabolic model^45^.

| Parameter | Name | Value |
| --- | --- | --- |
| $\delta$ | Gut dilution rate | 0.5 |
| $[G]_{\text{diet}}$ | Dietary glucose concentration | Varies |
| $[L]_{\text{diet}}$ | Dietary lactic acid concentration | Varies |
| $k_{a,G}$ | Glucose absorption rate | 100 |
| $k_{a,L}$ | Lactic acid absorption rate | 100 |
| $k_{\text{glycolysis}}$ | Glycolysis rate | 1.5 |
| $k_{\text{Ldh-f}}$ | Forward Ldh rate | 1 |
| $k_{\text{Ldh-r}}$ | Reverse Ldh rate | 0.01 |
| $k_{\text{OXPHOS}}$ | OXPHOS rate | 0.1, varies in regulation model |
| $k_{e,G}$ | Glucose elimination rate | 0.1 |
| $k_{e,L}$ | Lactic acid elimination rate | 0.1 |
| $k_{e,P}$ | Pyruvate elimination rate | 0.1 |
| $Y_{\text{glycolysis}}$ | Per-glucose ATP yield of glycolysis | 2 |
| $Y_{\text{OXPHOS}}$ | Per-pyruvate ATP yield of OXPHOS | 17 |
| $k_{\text{OXPHOS}}^{\text{max}}$ | Maximum OXPHOS rate | 10 |
| $f_{\text{basal}}$ | Basal fraction of OXPHOS activity | 0.01 |
| $r$ | Optimal ratio of glucose to lactic acid | 0.5 |
| $w$ | Width of OXPHOS regulation function | 0.13 |

## Notes

### Competing Interest Statement

The authors have declared no competing interest.

### Summary of Updates

The section on model parameterization has been updated to better align with formatting conventions. These do not change any results but aid clarity.

